# Paracrine enhancement of tumor cell proliferation provides indirect stroma-mediated chemoresistance via acceleration of tumor recovery between chemotherapy cycles

**DOI:** 10.1101/2023.02.07.527543

**Authors:** Daria Miroshnychenko, Tatiana Miti, Pragya Kumar, Anna Miller, Mark Laurie, Nathalia Giraldo, Marilyn M. Bui, Philipp M. Altrock, David Basanta, Andriy Marusyk

## Abstract

The ability of tumors to survive therapy reflects both cell-intrinsic and microenvironmental mechanisms. Across many cancers, including triple-negative breast cancer (TNBC), a high stroma/tumor ratio correlates with poor survival. In many contexts, this correlation can be explained by the direct reduction of therapy sensitivity by stroma-produced paracrine factors. We sought to explore whether this direct effect contributes to the link between stroma and poor responses to chemotherapies. Our *in vitro* studies with panels of TNBC cell line models and stromal isolates failed to detect a direct modulation of chemoresistance. At the same time, consistent with prior studies, we observed treatment-independent enhancement of tumor cell proliferation by fibroblast-produced secreted factors. Using spatial statistics analyses, we found that proximity to stroma is often associated with enhanced tumor cell proliferation *in vivo*. Based on these observations, we hypothesized an indirect link between stroma and chemoresistance, where stroma-augmented proliferation potentiates the recovery of residual tumors between chemotherapy cycles. To evaluate the feasibility of this hypothesis, we developed a spatial agent-based model of stroma impact on proliferation/death dynamics. The model was quantitatively parameterized using inferences from histological analyses and experimental studies. We found that the observed enhancement of tumor cell proliferation within stroma-proximal niches can enable tumors to avoid elimination over multiple chemotherapy cycles. Therefore, our study supports the existence of a novel, indirect mechanism of environment-mediated chemoresistance that might contribute to the negative correlation between stromal content and poor therapy outcomes.

## Introduction

For many types of cancers, standard cytotoxic chemotherapy remains the main treatment option. In many types of cancer, chemotherapies induce potent clinical responses that can translate to cures. On the other hand, despite having strong side effects, they often fail to eradicate cancers, especially in advanced, metastatic stages. Improving the clinical outcomes of these therapies requires an adequate understanding of the mechanisms responsible for treatment failure. These needs have motivated massive research efforts to identify a large and growing list of specific molecular mechanisms responsible for chemoresistance. Whereas chemoresistance studies have primarily focused on cell-intrinsic genetic and epigenetic mechanisms, there is a growing appreciation of the importance of cell-extrinsic, microenvironmental factors. Across many types of cancers, high stroma-to-tumor ratios correlate with poor outcomes (1–8). In multiple contexts of targeted therapy, this correlation appears to reflect the effects of direct microenvironmental resistance mechanisms. Multiple growth factors and extracellular matrix (ECM) components produced by cancer-associated fibroblasts (CAFs) and other non-malignant cell types within the tumor microenvironment (TME), can strongly blunt the sensitivity of cancer cells to inhibition of oncogenic drivers by the activation of alternative signaling pathways and the induction of cell plasticity (9–11). However, the relevance of direct desensitization toward chemotherapy sensitivity is less clear. We sought to explore the link between stroma and chemotherapy responses in triple-negative breast cancers (TNBC), where stromal gene expression signature is linked to poor chemotherapy responses (12). Despite developments in immune therapies and the recent approval of PARP inhibitors and antibody-drug conjugates, cytotoxic chemotherapies remain the primary therapeutic tool for adjuvant and neoadjuvant TNBC therapy. The most commonly used treatment modality in the US is a dose-dense therapy that includes <u>a</u>nthracyclines (such as doxorubicin), <u>c</u>yclophosphamide, and taxanes (AC-T regimen).

Chemotherapy is administered intravenously as an injection over a few minutes or as infusions over a few hours. Two to three-week breaks are given to allow patients to recover from the strong side effects of the treatment. This cycle is repeated several times. TNBC tumors are typically sensitive to chemotherapy, and this sensitivity is maintained over the course of the treatment (13), suggesting that TNBC can avoid therapeutic elimination without acquiring *bona fide* resistance (14) as long as a sufficient number of tumor cells can survive till the end of chemotherapy.

Like several other cancer types, high stromal content in TNBC strongly correlates with poor prognosis (7,12,15). Moreover, stromal gene expression signatures predict a diminished response to the anthracycline-containing chemotherapeutic regimen in neoadjuvant therapy (12), suggesting a causal link between stroma and therapy persistence. However, we failed to observe direct chemoprotection with a panel of CAF isolates and TNBC models. Instead, in agreement with prior studies documenting the growth-promoting effects of CAFs (9,16,17) we found that, often, CAFs and CAF-conditioned media (CAF CM) enhance the proliferation of TNBC cells *in vitro*, independent of the treatment. Consistent with these observations, we found that proximity to stroma is associated with enhanced tumor cell proliferation *in vivo*, in xenograft models and clinical samples. These findings led us to hypothesize an indirect link between stroma and chemoresistance: the ability of tumors to avoid therapy-induced extinction is mediated by an accelerated recovery of residual tumors between cycles of chemotherapy. To test the feasibility of this hypothesis, we developed a mathematical model (a spatial agent-based model (ABM)), parameterized and calibrated with our experimental data, to understand the impact of stroma-enhanced proliferation on the cancer cell population dynamics. We found that a relatively modest enhancement of cell proliferation within stroma-proximal tumor cells is sufficient to substantially decrease the probability of therapeutic extinction. Our findings indicate the existence of a new type of indirect chemoresistance mechanism that, acting in parallel with cell-intrinsic chemoresistance mechanisms, likely contributes to the ability of a subset of tumors to persist through therapy and might be suppressed therapeutically.

## Results

### CAFs stimulate the proliferation of TNBC cells but provide no direct chemoprotection

CAFs are one of the main cellular components of the TME and the primary producer of ECM, growth factors, and cytokines. Multiple cytokines and ECM components, produced by CAFs (e.g., HGF, FGFs, fibronectin, *etc*.) have been shown to strongly reduce the sensitivity of cancer cells to therapeutic agents in multiple contexts of targeted and cytotoxic therapies (6,18–21). Therefore, we asked whether the known correlation between poor prognosis and high stromal gene signature can be reduced to a correlation with CAFs. To this end, using the KM plotter tool (22) (see Methods), we interrogated publicly available expression datasets to explore the association between the expression of CAF markers and TNBC survival. We found that the expression of individual CAF markers *α*SMA, FAP, and PDGFRa, was associated with significantly lower survival (**Fig. S1**). This association led us to speculate that, like other contexts of CAF-mediated therapy resistance (23,24), CAFs might directly reduce the sensitivity of TNBC cells to chemotherapeutic agents. To test this hypothesis, we evaluated the ability of CAFs to modify the sensitivity of a panel of TNBC cell lines to the AC-T components doxorubicin and taxol, as well as the widely used therapeutic agent cisplatin. Since the commonly used viability assays, such as CellTiterGlo or MTT cannot discriminate between the viability signals of cancer cells and CAFs, we used luciferase-labeled TNBC cells and bioluminescence readout, as described in our prior work (25) (**Fig. 1A**). When normalized to vehicle-treated TNBC cell monocultures (**Fig. 1B**), co-culture with CAFs enhanced the luminescence readout under treatment in most of the tested models (**Fig. 1C**). However, we also observed a substantially higher luciferase signal in the DMSO controls of the CAF co-cultures, suggesting a general enhancement of proliferation, rather than reduction of sensitivity to therapy. Indeed, when the readouts of the CAF co-cultures were normalized separately to their DMSO controls (**Fig. 1B**), luciferase signal enhancement under therapy was no longer observed (**Fig 1D**). Notably, this lack of specific protection stands in contrast to the CAF-mediated desensitization to the therapies that target specific signaling nodes, such as EGFR/HER2 inhibitor lapatinib, reported in our prior work (25), or pan-class I PI3K inhibitor BKM120 (**Fig. S2A)**.

**Figure 1.**
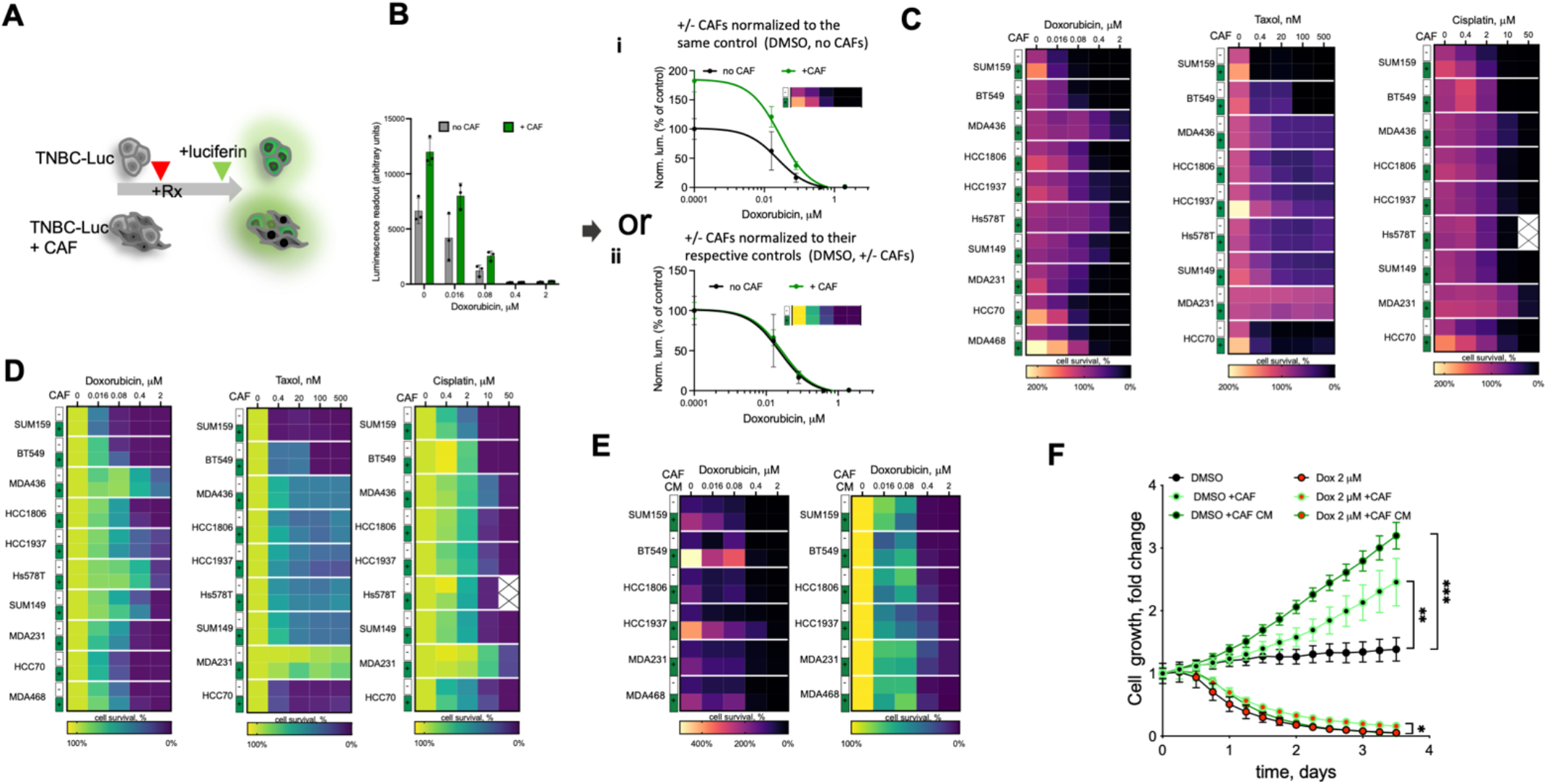
CAFs facilitate TNBC proliferation *in vitro*. **A**. Experiment diagram for the chemosensitivity sensitivity assay. Luciferase-labeled TNBC cells are cultured in the presence or absence of unlabeled CAFs in the presence of doxorubicin or DMSO vehicle control. Only TNBC cells directly contribute to the viability signal. **B**. Normalization schemata for the data analyses. Raw data from the viability assay can be normalized to either i) the DMSO control signal of cells cultured without CAFs, or ii) with separate normalization of the control and CAF co-cultures to their respective DMSO controls. **C, D**. Heatmap summaries of the impact of CAF co-cultures on the sensitivity of the indicated chemotherapeutic agent in a panel of TNBC cell lines, normalized as i) or ii) in panel **B**, respectively. **E**. Heatmap summaries of the impact of CAF CM on doxorubicin sensitivities of the indicated TNBC cell lines. **F**. Impact of CAFs and CAF CM on the growth of GFP-labelled MDA468 cells following 24 hours of doxorubicin exposure, measured by time-lapse microscopy. Statistical analyses of indicated differences were performed with a paired 2-tail t-test, comparing confluency value at each of the time point *** p=0.0007, ** p=0.003, * p=0.0102.

A general enhancement of the luciferase signal was also observed with a large panel of primary human CAFs, normal breast fibroblasts, and immortalized and primary mouse fibroblasts (**Fig. S2B, C**). Of note, consistent with the prior knowledge, CAFs were much less sensitive to chemotherapy, compared to TNBC cells (**Fig. S2D**).

The effect of CAFs was phenocopied with CAF-conditioned media (CM), suggesting mediation by secreted factors (**Fig 1E**). To establish whether higher luciferase readouts reflected growth-promoting or metabolic effects of CAFs, we assessed cell proliferation more directly with time-lapse microscopy using TNBC cells labeled with nuclear GFP, as described in our prior work (26). To facilitate the ability to detect an enhancement of cell expansion, cells were seeded at low (∼10%) confluency. CAFs co-culture and CAF CM potently enhanced proliferation in multiple TNBC cell models (**Fig. 1F** and **S2E**). In contrast, while CAFs and CAF CM enhanced viability under doxorubicin treatment, the effect was much weaker than in the DMSO controls (**Fig. 1F**), suggesting that this enhanced viability reflects a general enhancement of TNBC proliferation rather than a direct enhancement of chemoresistance. Finally, the proliferation-enhancing effect of CAFs and CAF CM was not limited to 2D cultures, as a similar effect could be observed within 3D cultures (**Fig. S2F, G**).

### Proximity to stroma enhances TNBC cell proliferation *in vivo*

In contrast to the well-documented ability of CAFs to enhance tumor cell proliferation in vitro, the existence of this effect *in vivo* is less clear. While the ability of CAFs to promote tumor growth is well-established, this does not necessarily imply enhanced tumor cell proliferation, as cell proliferation is not strictly coupled with tumor growth (27). Moreover, CAFs are also known to impact additional phenotypes that can limit tumor growth, including neovascularization, immune surveillance, as well as the ability of tumor cells to migrate and invade the surrounding tissues (9). Therefore, we sought to assess the *in vivo* relevance of the enhanced proliferation observed *in vitro* more directly. The establishment and growth of experimental tumors in murine xenograft models are contingent on the recruitment of host stroma. CAFs play an essential “infrastructural” role by producing ECM that provides the structural organization of tumor tissues, e.g., enabling cell-ECM adhesion and vascularization, *etc*. Therefore, in contrast to *in vitro* studies, a “clean” +/-CAF comparison in experimental tumors is impossible. At the same time, even genetically homogeneous experimental tumors typically display substantial spatial heterogeneity, including regional differences in the stroma-to-tumor ratios. We leveraged this variability by assessing the correlation between stromal content and tumor cell proliferation across semi-randomly selected tumor microdomains of variable tumor tissue across whole tumor cross-sections (**Fig. 2A**). To enable accurate detection of proliferating cells, animals bearing experimental tumors were injected with the dNTP analog BrdU before the euthanasia. Following IHC staining against BrdU, images of whole-slide scans were segmented into micronecrotic regions, stroma, BrdU+ & BrdU-tumor cells using the AI-assisted digital pathology platform Aiforia (https://www.aiforia.com)(**Fig. 2A, S3A**). Consistent with our *in vitro* findings, we found a strong correlation between the fraction of BrdU+ tumor cells and stromal content in MDA468 and BT549 xenograft models of TNBC, under both vehicle control and doxorubicin or taxol administration (**Fig. 2B, S3B**).

**Figure 2.**
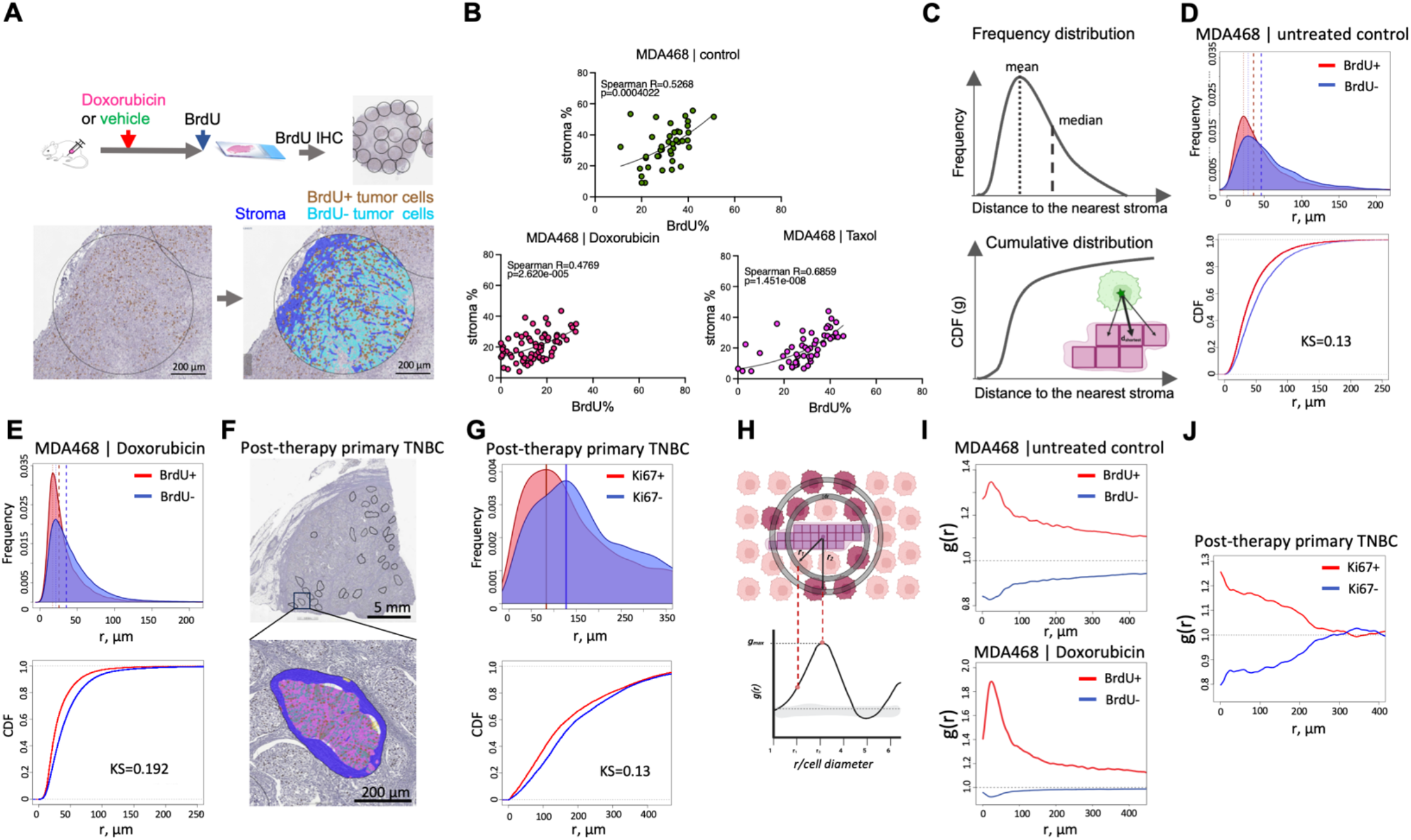
Proximity to stroma correlates with higher proliferation *in vivo*. **A**. Diagram of the experimental approach to assess the impact of stroma proximity on the proliferation of TNBC cells *in vivo*. Before euthanasia, the mice are pulsed with BrdU, which enables IHC-based detection of cells in the S phase of the cell cycle. Tumor tissue in whole slide scans of BrdU IHC staining is subsampled into smaller areas (0.9 mm in diameter); necrosis-free tumor tissue within these subsampled regions is segmented into BrdU+/-tumor cells and stroma. **B**. Regression analyses of MDA468 xenograft tumors, treated with doxorubicin (2.5 mg/kg), Taxol (10 mg/kg) or vehicle control 48 hours before euthanasia were used to assess the correlation between stromal content and tumor cell proliferation. Each dot represents a subsampled ROI, as in A. Spearman R and p values of non-linear fit are shown. **C**. Schemata for the nearest neighbor analyses that calculate distances between each of BrdU+/-cells in the tumor cross-section and the nearest stromal pixel. **D, E**. Frequency distribution and cumulative distribution function (CDF) plots of distances to the nearest stroma of BrdU+/-cells in MDA468 xenografts tumors from control (**D**) and doxorubicin-treated (**E**) mice. Dashed lines indicate medians and means of the distributions KS denotes the Kolmogorov-Smirnov statistical test. **F**. A representative image of a diagnostic biopsy of a post-treatment primary human TNBC tumor, stained with proliferation marker Ki67, ROIs used for subsampling, and an example segmentation of an ROI into Ki67+/-cells and stroma. **G**. Frequency distribution and CDF plots of distances of Ki67+/-cells to the nearest stroma in the primary TNBC sample. **H** Schemata for the RDF analysis. **I**. RDF analyses of the impact of stroma proximity on the distribution of BrdU+/-cells in control and doxorubicin-treated MDA468 tumors. **J**. RDF analyses of the impact of stroma proximity on the distribution of Ki67+/-cells in a post-treatment primary TNBC tumor.

Encouraged by these observations, we sought to validate these findings with an independent approach that relies on analyses of the distributions of distances between BrdU+ and BrdU-cells and the nearest stroma (**Fig. 2C**). In these analyses, every tumor cell within the tumor cross-section was analyzed, thereby minimizing bias, and improving statistical power. Consistent with the correlation-based analyses, we found a significant difference in the distribution of BrdU+ and BrdU-cells, with BrdU+ cells distributed, on average, closer to the stroma, supporting the notion that stroma-produced paracrine factors stimulate cell proliferation *in vivo* (**Fig. 2D, E**). Importantly, our analyses revealed a similar bias in the distribution of Ki67+ cells in a post-therapy sample of primary TNBC, suggesting broader relevance to primary human disease (**Fig. 2F, G**).

To quantitatively assess the impact of stroma on the proliferation of tumor cells *in vivo*, we decided to repurpose a radial distribution function (RDF), **g**(**r**), which is often used in the field of spatial ecology (28–30). RDF can be used to compare the average density of observed BrdU+/-tumor cells against randomly distributed BrdU+/-across different length scales from each of the stromal pixels. The location of the RDF peak, **g**_**max**_, indicates the linear distance of the effect of stroma on cell proliferation. The **g**_**max**_ value indicates the magnitude of the effect (**Fig. 2H**). To improve the accuracy of our analyses, our analyses were applied to whole histological cross-sections. Thus, each analysis captured the spatial patterns of BrdU positivity of 10^4^ to 10^6^ cancer cells per tumor, with multiple tumors analyzed per experimental condition.

The RDF revealed that, in MDA468 xenograft tumors, the proliferation probability of cells within 2-3 cell diameters from the stromal boundary (∼ 30 μM) was increased by 30-40% in the chemotherapy naïve tumors and further increased after doxorubicin treatment (**Fig. 2I**). A similar (25-30%) enhancement was also observed in a primary post therapy TNBC sample, with the effect distributed over a larger distance from stromal edge (**Fig. 3J**). In summary, these analyses support the notion of the in vivo relevance of proliferation-enhancing effects of stroma, while also highlighting the spatial limitation of this effect.

**Figure 3.**
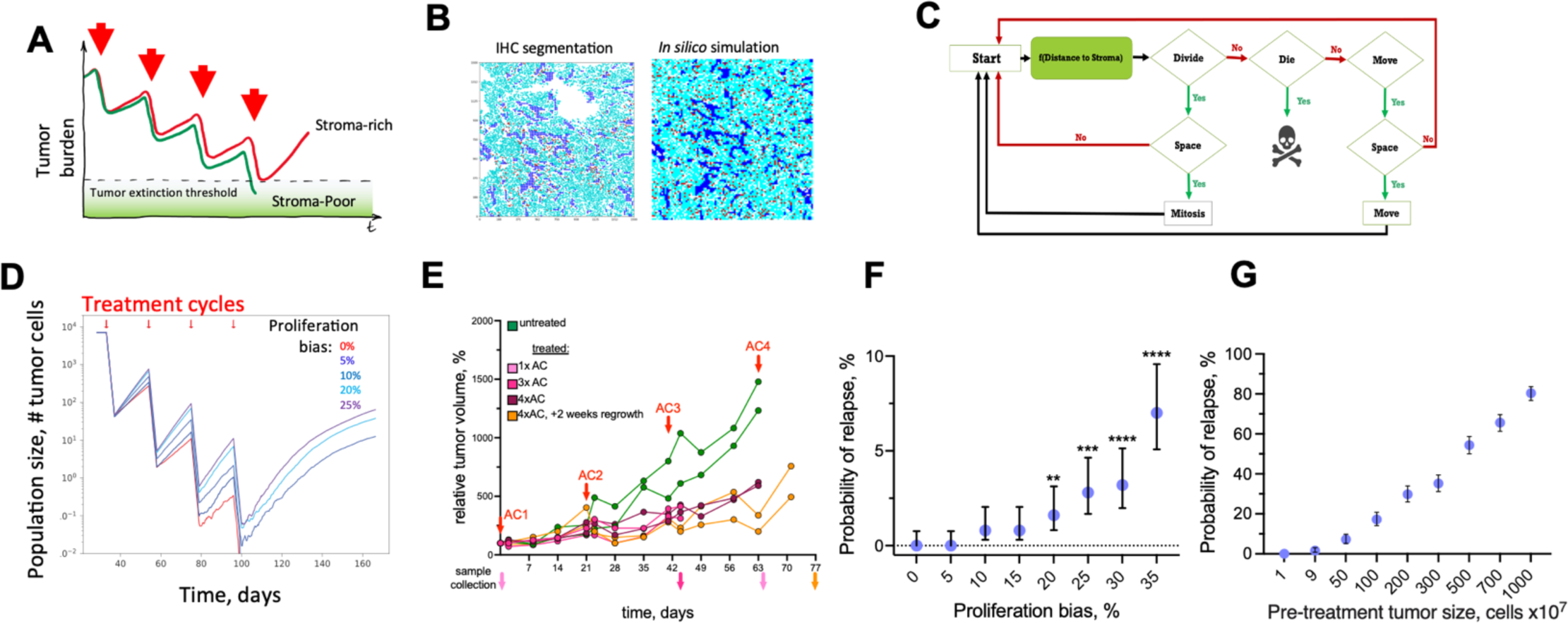
In silico validation of the hypothesized indirect stroma-mediated chemoresistance. **A**. Model schemata depicting the hypothesized indirect stroma-mediated chemoresistance. Enhanced proliferation in stroma-rich tumors can enhance between-chemo cycles recovery of tumors, enabling them to escape therapeutic eradication. **B**. ABM is initiated based on the spatial localization of tumor cells and stroma observed in the indicated experimental sample. **C**. Diagram of the ABM model. **D**. Dynamics of volume changes in MDA468 xenograft tumors over the course of AC treatment (0.5 mg/kg doxorubicin and 50 mg/kg cyclophosphamide), injections times are indicated by red arrows. Traces indicate individual tumors; distinct colors of volume traces indicate tumors harvested at different time points, indicated by arrows of matching colors. **E**. Impact of the indicated magnitude of enhancement of cell proliferation within 3 cell diameters from stroma border on the average population size over the course of chemotherapy. Traces depict average population sizes over 500 simulations per condition. **F**. Impact of the indicated magnitude of enhancement of cell proliferation on the probability of tumor relapse through the course of therapy, over 500 simulations with 95% confidence interval. ****, ***, ** indicate p-values of <0.001, <0.001, and 0.0076, respectively, of Fisher exact test, comparing the probability of relapse with indicated proliferation bias against the simulations without stromal effect (proliferation bias 0%). **G**. Dependence on the sampling grid size of the tumor relapse for the simulations under short-term cytotoxic effects of the chemotherapy under 5% bias in proliferation due to stromal effects. For each data point, 500 random samplings of groups of 1,9,50, 100, 200, 300, 500 700, and 1000 simulations have been randomly selected from 10,000 simulations of 100×100 grids. Error bars depict 95% confidence intervals.

### *In silico* model for characterization of proliferation/death dynamics in time and space

Whereas the ability of fibroblasts to enhance tumor cell proliferation has been observed in multiple contexts, it is unclear whether the enhanced proliferation can blunt the effects of chemotherapy. Higher proliferation rates are considered to enhance rather than decrease sensitivity to DNA-damaging cytotoxic chemotherapy (31). (Therefore, in the absence of direct stroma-mediated resistance, one would expect the proliferation-enhancing effects of stroma to increase rather than decrease chemoresistance. On the other hand, the differences in chemosensitivity are typically assessed within short time scales that do not necessarily reflect tumor response over the course of chemotherapy treatment. Cytotoxic chemotherapies, including the ACT regimen used in TNBC therapy, are administered in several cycles of brief chemotherapy exposures separated by several weeks of recovery (32). This pattern administration is based on the well-established concept of a fractional cell kill, *i.e*., chemotherapy exposure eliminates a fraction of the neoplastic population with each cycle (33), thus necessitating repeated dosing to achieve complete elimination. The length of the recovery between administrations of chemotherapy is typically designed with the aim of minimizing systemic toxicity. Given the short duration of conventional drug administration and the short half-life of chemotherapeutic agents, the surviving tumor cells have an opportunity to partially recover, re-populating the tumor between therapy cycles. Cancer is cured if, over the course of multiple cycles, all the tumor cells are eliminated or if the surviving fraction is below a threshold for stochastic or immune-mediated extinction. Thus, we hypothesized that the enhanced proliferation of stroma-proximal cells could reduce chemosensitivity by enhancing the ability of residual tumor cell populations to regrow between therapy cycles (**Fig. 3A**). We expect this effect to contribute to other factors shaping chemotherapy responses, including cell-intrinsic drug sensitivity, dormancy, inequality of drug access, selection for cells with reduced sensitivity over multiple cycles, direct environmental chemoresistance, *etc*. Thus, from the first principles, the effect of enhanced proliferation should be most impactful when therapy eliminates the majority, but not all, of tumor cells at each cycle. For intrinsically chemoresistant tumors, the effect is expected to have a marginal contribution. Likewise, the hypothesized effect would be irrelevant if all the tumor cells were eliminated or arrested for the whole duration of individual chemotherapy cycles.

We sought to evaluate the potential relevance of this hypothesis under a biologically relevant set of conditions. Evaluating this hypothesis requires simultaneous consideration of the combination of factors, including the initial size of the neoplastic population, proliferation/death rates of tumor cells during both acute and recovery phases of chemotherapy cycles, the magnitude of proliferation enhancement effect, and the size of the peristomal niche. Given the lack of experimental *in vivo* models enabling to turn the effects of stromal niche on and off, separate the effect of stromal niche from the additional mechanisms that influence chemoresistance, and achieve strong, but incomplete responses, “clean” evaluation of the hypothesis presented in **Fig. 4A** necessitated an *in-silico* examination. To this end, we constructed an on-lattice stochastic, agent-based model (ABM) capable of integrating the spatiotemporal effects of stroma on proliferation and death dynamics using the HAL platform (34) (see Supplemental information for full details). The ABM is initialized as a 100×100 cell grid with the location of tumor cells and stroma matching those observed within a region from a histological sample (**Fig. 3B)**. In our model, tumor cells are “agents”, while stroma remains static throughout the simulation. At every time step of the simulation, each of the tumor cells can either remain unchanged, die, divide, or move to an adjacent space on the lattice, with division and movement dependent on space availability. Proliferation and death probabilities are influenced by the effects of chemotherapy and proximity to stroma (**Fig. 3C)**.

**Figure 4.**
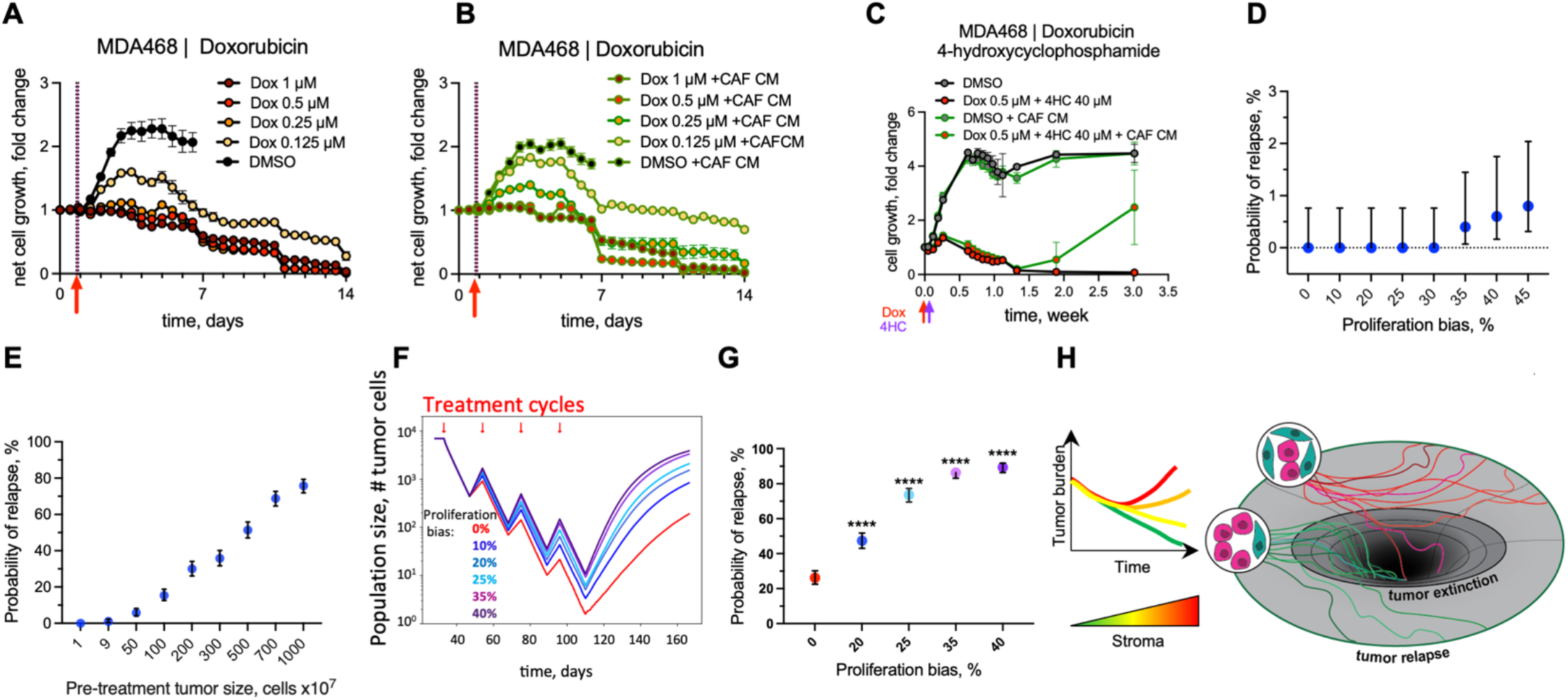
Impact of the stroma-enhanced proliferation on tumor chemotherapy recovery. **A, B**. Changes in numbers of viable MDA468 cells following brief (2-hours) administration of doxorubicin in control medium (**A**) and with CAF CM (**B**). **C**. Impact of CAF CM on the response of MDA468 to the sequential administration of doxorubicin and 4-hydroxy cyclophosphamide (1 hour each). **D**. Probability of relapse for the simulations under long-term cytotoxic effects of the chemotherapy, with the killing fraction of 21% of the cells per time step. Each data point in the graph represents the average of 500 simulations, error bars indicate 95% confidence interval. **E**. Dependence of the relapse probability on the initial population size under the scenario of 20% bias in tumor cells proximal to the stroma. Each data point is the average of the 500 samplings and the respective 95% confidence interval. **F**. Impact of the indicated magnitude of enhancement of cell proliferation within 3 cell diameters from stroma border on the average population size over the course of chemotherapy that, in the absence of stromal effects, eliminates tumors with 27% relapse probability. Traces depict average population size over 500 simulations per condition. **G**. Probability of tumor relapse affected by different proliferation bias and prolongated cytotoxic effect. Graphs depict the outcomes of 500 simulations with 95% confidence interval. **** indicates < 0.001 p-values of Fisher exact test, comparing outcomes with different values of stromal enhancement values to simulations without stromal effect (proliferation bias 0%). **H**. Conceptual model of the indirect stromal chemoresistance. The link between high stromal content and chemoresistance might be at least partially mediated by the stroma-dependent potentiation of tumor cell proliferation, which enhances tumor recovery between chemotherapy cycles and decreases the odds of chemotherapeutic extinction.

The utility of *in silico* simulations depends not only on the validity of the model’s assumptions but also on the quantitative accuracy of the parameters that define the dynamics of agents within the model. Even though therapy outcomes are shaped by multiple cell-intrinsic and cell-extrinsic mechanisms, the effects of these mechanisms ultimately converge on the balance between cell proliferation and cell death. Despite the fundamental importance of these parameters, their reliable estimates are lacking in the literature, forcing mathematical modelers to operate with net growth rates, which can be inferred from experimental and clinical data with relative ease and accuracy. However, the same net growth rate can result from highly different birth (proliferation) and death rate combinations. For example, while stagnant tumor growth can be a manifestation of proliferation rate, it can also result from space restriction or other resource limitations, resulting in high turnover rates, with high rates of cell proliferation being balanced by high levels of cell death (27). Xenograft models enable accurate estimation of proliferation rates owing to the availability of experimental labels like BrdU. BrdU, which is a dTTP analog, incorporates into DNA during the replication (S) phase of the cell cycle (35). Thus, a brief BrdU label administered prior to tumor harvest provides an accurate quantitative estimate of the fraction of cells in the S phase of the cell cycle. Since the duration of the S phase in mammalian cells is highly conserved (8-12 hours), the proportion of BrdU+ cells can be used to accurately quantify tumor birth rates (36). Whereas similar direct estimates of cell death rates are not available, they can be inferred as a difference between cell proliferation, inferred from BrdU labeling, and net changes in disease burden available from volumetric studies. Given the inability of common xenograft models to accurately recapitulate clinical chemotherapy sensitivity, we decided to use a hybrid approach, where proliferation rates are directly estimated from xenograft studies while death rates are determined from the death rates, sufficient to eradicate the tumors over the course of multiple therapy cycles in the absence of stromal enhancement of proliferation.

To infer cell proliferation during therapy, we subjected a cohort of mice bearing MDA468 xenograft tumors to a clinically relevant regimen of four cycles of AC (doxorubicin/cyclophosphamide) treatment with three-week intervals between cycles. Tumors were harvested two days after the first, third, and fourth cycles, with additional 2 tumors harvested two weeks after the fourth cycle. Consistent with generally weak chemotherapy responses in xenograft TNBC models, AC treatment slowed the growth of MDA468 xenograft tumors but failed to induce substantial tumor regression (**Fig. 3 D**). Similar to the observed responses of MDA468 xenografts to a single dose of doxorubicin **(Fig. S2B)**, the doxorubicin/cyclophosphamide combination led to a ∼ 2-fold reduction in the fraction of BrdU+ cells, and this suppression lasted over the subsequent AC cycles (**Fig. S4A**). Strong cytotoxic responses are expected to increase the proportion of stroma due to stroma activation and elimination of tumor cells. However, consistent with the lack of strong response to the chemotherapy, we observed only a slight increase in the stroma-to-tumor ratio (**Fig. S4B**), while the effect of therapy on the fraction of stromal area and the proportion of cells staining positive for the apoptotic marker cleaved caspase 3 did not reach statistical significance (**Fig. S4D**). To estimate the effects of therapy on cell proliferation and tumor/stroma ratios, we analyzed six pairs of matched pre- and post-therapy histological samples in TNBC patients treated with a full course of AC-T neoadjuvant therapy without achieving full pathological response. As expected, post-therapy samples displayed a trend to increase in the stroma area, with reduced cellularity and cell proliferation (**Fig. S5A, B**). The effects were not statistically significant, likely reflecting substantial patient-patient differences, small sample size, and variability in the time between the AC-T completion and surgery. However, analysis of a larger histological dataset of unpaired samples revealed a statistically significant increase in the stroma, as well as a bimodal distribution of the effect of therapy on proliferation (**Fig. S5C**).

Given that the MDA468 model adequately recapitulates clinically relevant baseline growth and proliferation proliferative index (27)and that the reduction in proliferation rates in xenograft tumors is consistent with therapy responses in at least some human tumors, we reasoned that the xenograft data is suitable for parameterization of cell proliferation rates during therapy. Therefore, we assumed that tumors maintain constant proliferation during therapy (with a rate reduced two-fold compared to pre-therapy), with therapy inducing a brief period of substantially enhanced death, followed by the return of death rates to the baseline during the recovery periods before the subsequent therapy cycle. Following these considerations, from the BrdU labeling data, we estimated the average proliferation rate of 0.198 per day during therapy. The knowledge of proliferation rates and net growth rates enabled us to estimate the death rate outside of the acute phase of therapy to be 0.165 per day (see Supplementary Methods).

Next, we used the ABM simulation to determine the acute chemotherapy-induced death that, in the absence of the stromal enhancement, is sufficient to eliminate *in silico* tumors over the course of four chemotherapy cycles under the fractional cell kill scenario. Given the short infusion duration and the rapid pharmacokinetics of chemotherapeutic agents *in vivo*(37), the acute therapy-induced DNA damage that acts as a trigger of cell death should be inflicted within hours. Cell viability assays (such as the MTT or Cell Titer Glo), commonly used in preclinical *in vitro* studies to evaluate therapeutic sensitivities, typically measure cell viability after three to four days post-drug administration. Therefore, in our *in silico* simulations, we decided to equally distribute acute therapy-induced tumor cell death over the course of four days (without directly affecting cell birth rates), at which point cell death rates to the baseline. An example of a simulation run of the model is shown in **video S1**. We found the minimal therapy-induced death rate that is sufficient to eliminate *in silico* tumors with complete penetrance (100% extinction rate in 500 simulations) after four chemotherapy cycles in the absence of stroma proximity to be 0.45 per day (**Fig. S6 A, B**).

### Testing the impact of stroma-enhanced proliferation on tumor eradication

The development and parameterization of the *in silico* model enabled us to assess the impact of enhanced proliferation within stroma-proximal niches on the odds of therapeutic extinction. To this end, we asked how the enhancement of proliferation within 3 cell diameters from stroma impacts the outcomes of treatment that is otherwise sufficient to achieve a full penetrance (in 500 simulations) over the course of treatment. We found that a 10% or higher enhancement of proliferation enabled some of the *in silico* tumors escape eradication, with the higher proliferation enhancement increasing the probability of escape. (**Fig. 3 E, F, Video S2**). An increase in the initial population size enhanced the probability of relapse. Even under a 5% proliferation bias which was insufficient to mediate rescue when the initial population was below 1×10^4^ cells, enabled 80% relapse when the initial population size was 1×10^7^ cells (**Fig. 3G**); the effect was even more pronounced with a 15% enhancement (**Fig. S6C**). Thus, the slight enhancement of cell proliferation, restricted to stroma-proximal cells, might enable an ecological rescue even in the absence of cell-intrinsic chemoresistance.

Next, we sought to assess whether this rescue effect can be recapitulated experientially. Given the lack of appropriate *in vivo* models, we decided to assess the effect *in vitro*, where proliferation-enhancement effects of stroma could be recapitulated by the addition of CAF CM. To recapitulate the scenario of high-intensity, transient clinical exposure, chemotherapy was applied for 2 hours, then the media was replaced with a drug-free media after a thorough PBS wash. Based on the commonly used 3-4 duration of drug dose-response assays, we expected that, under this brief exposure scenario, the cells that survive up to day 4, will be able to recover and regrow. Surprisingly, we found that the cytotoxic impact of the brief doxorubicin exposure on the viability of tumor cells extended over a substantially longer time frame and that even substantially lower doxorubicin concentrations allowing for an initial expansion, led to the continuous decline of viable tumor cells for at least 2 weeks post the brief exposure (**Fig. 4A**).

Notably, the decline in cell numbers occurred despite the active proliferation of tumor cells, consistent with a lasting induction of increased cell death rates (**Video S3**). The longer-lasting effect of chemotherapy was also observed in additional models of TNBC that we have tested (HCC1806 and HCC1937), although the duration of the effect and the baseline doxorubicin sensitivities varied with the model (**Fig. S7A, B**). Similar lasting suppression of net proliferation was observed with taxol (**Fig S7C**). Moreover, the lasting effect of chemotherapy was observed with A678 cell line model of Ewing sarcoma, following exposure to doxorubicin, vincristine, and 4-HU (**Fig. S6C**), suggesting a broader generalizability of the lasting chemotherapy impact towards different cancer types and chemotherapeutic agents. Finally, the extended effect was not limited to 2D studies, as treatment with doxorubicin and doxorubicin - 4HC (active metabolite of cyclophosphamide) combination induced loss of viability that lasted 7-10 days (**Fig. S7E, F**)

The addition of CAF CM (**Fig. 4B, Fig. S7A-C, F**) or physical co-culture with CAFs (**Fig. S7E**) partially counteracted the lasting cell attrition induced by transient chemotherapy administration across all of the tested TNBC models. While automatic cell detection had insufficient sensitivity to detect rare viable cells, visual examination of the time-lapse sequences indicated that under conditions that led to complete elimination of controls (**Video S3**), viable cells could often be detected in cultures with CAF CM (for example, see **VideoS3, S4** for doxorubicin treated MDA468 cells). Notably, we also observed a more obvious rescue in MDA468 cells, treated with sequential 1 hr exposure of doxorubicin and 4HC (**Fig, 4C**).

Whereas the in vitro studies provide proof of principle validation of our hypothesis, presented in **Fig. 3A**, in vitro studies cannot recapitulate the in vivo birth/death dynamics. For example, CAF CM mediated rescue shown in **Fig. 4C** enabled surviving cells to expand above the initial cell numbers. On the other hand, the lingering cytotoxic effects of therapy are expected to reduce the impact of the postulated effect of stroma *in vivo*. Thus, we re-evaluated our *in silico* simulations, taking the extended duration of chemotherapy induced cell death into account. While we observed a substantial variability in the duration of the lingering therapy-induced death, we decided to use a more conservative two-week estimate, as the shorter recovery time should limit the effect of the accelerated recovery.

We started by assessing whether the brief chemotherapy exposure leads to a single or bi-phasic (acute phase followed by lowering intensity lingering effects) induction of cell death. The fitting of *in vitro* experimental data indicated that chemotherapy induces a monotonous decrease in cancer cell numbers, consistent with a single phase (**Fig. S8A &** Mathematical Supplement). Next, after finding the threshold death rate that sufficient to eliminate tumors in all 500 *in silico* tumors, we asked whether the enhanced proliferation within stroma proximity can still counteract elimination. The extended duration of chemotherapy-induced cell death translated into a reduction in per-day death rates, which is necessary to eliminate the tumors over the course of four cycles of chemotherapy from 0.45/day (**Fig. S6A, B**, to 0.2/day (**Fig. S8B, C**). As expected, the reduced recovery time counteracted the ability of stroma to rescue tumors from therapeutic extinction but did not entirely negate this effect. The threshold proliferation–enhancing effect of stroma, detectable with 500 simulations, increased from 10% under the four-day duration of tumor cell kill (**Fig. 3 H, I**) to 35% under the two-week duration (**Fig. 4D, S8D)**. Increasing the number of simulations to 10,000 shifted the threshold to 20%; similar to the 4-day acute effect duration, an increase in the initial population size magnified the effect of stroma, shifting relapse probability to ∼80% when the initial population was 1×10^7^ cells (**Fig. 4E**). Next, we evaluated the impact of stromal enhancement of proliferation under lower therapy-induced death rates where, in the absence of stromal effects, therapy leads to a partial (75%) penetrance of tumor eradication under initial population under 1×10^4^ cells. Under this scenario, even a 10% enhancement of proliferation within 3 cell distances from the stroma almost doubled the relapse probability (**Fig. 4E, F**). The effect was even more pronounced with the stronger stromal enhancement (**Fig. 4E, F**).

In summary, our results indicate the existence of an indirect, stroma-mediated chemoresistance mechanism where enhanced proliferation within stroma proximal niches shifts the net outcomes of proliferation-death dynamics, enabling tumors at the brink of elimination to avoid therapy-induced extinction (**Fig. 3A and 4G**). More generally, our study highlights the importance of consideration of extending the consideration of causes of therapy resistance beyond proximal molecular mechanisms.

## Discussion

Despite the advances in basic and clinical cancer research, the combination of surgery and chemotherapy usually fails to eradicate advanced cancers, including TNBC. Improving clinical outcomes requires an accurate and comprehensive understanding of the biological mechanisms that enable tumors to avoid therapeutic elimination. Chemoresistance and patient-patient variability in clinical responses reflect a combined output of multiple contributing factors, including mechanisms acting on cell-intrinsic, microenvironmental, and systemic scales. The stochastic nature of somatic evolution, underpinning carcinogenesis, combined with genetic, environmental, and other differences between the cancer patients, leads to a substantial tumor-tumors variability in chemosensitivity. Further, intra-tumor genetic and epigenetic variability, as well as mutational and plasticity-mediated diversification, can lead to differential survival of chemoresistant subpopulations and declined responses over multiple cycles of chemotherapy. At the same time, cancers can also persist through chemotherapy without developing resistance, as the ability of some tumor cells to enter dormancy before or during therapy enables them to “wither the storm”, re-awakening to resume proliferation after therapy resumption (14,38). Our studies uncover a novel, indirect mechanism of stroma-mediated chemoresistance where treatment-independent enhancement of tumor cell proliferation in the stroma’s vicinity augments residual tumors’ ability to recover between chemotherapy cycles, thus decreasing the odds of therapeutic extinction over the course of treatment (**Figs. 3A)**. The magnitude of this mechanism’s impact on tumor eradication depends on the combined output of several parameters, including disease burden, proliferation/death rates, their modification by therapy (including the duration of therapy-induced changes), the spatial distribution of tumor cells and stroma, as well as the size and distance of the effects of stroma proximity. With all things being equal, under biologically relevant proliferation, death, and stromal effect parameters, the proliferation-enhancing impact of stroma can tilt the balance, enabling tumors to avoid therapeutic elimination (**Fig. 4G**). The proposed mechanism should be most impactful for those tumors on the brink of therapeutic extinction, *i.e*., when, in the absence of the effect, therapy achieves a partial penetrance of disease elimination.

Most of the research and development efforts in therapy resistance are directed to the identification and targeting of individual molecular mechanisms that enable tumor cells to avoid therapeutic elimination. This approach is not limited to cell-intrinsic mechanisms but also extends toward cell-extrinsic, microenvironmental mechanisms of therapy resistance. In the context of targeted therapies directed against oncogenic signaling pathways, multiple paracrine factors, including elements of the ECM and cytokines produced by CAFs and other non-neoplastic cells within the TME, have been documented to confer resistance to targeted inhibitors by activating alternative signaling pathways (39–41). A strong correlation between high stromal content, chemoresistance, and poor patient survival in TNBC (12), where cytotoxic therapy remains the main therapeutic option, suggests the existence of similar direct chemoresistance mechanisms. Indeed, several studies have reported direct stroma-mediated chemoresistance in the context of chemotherapy (42,43). However, the magnitude of the direct effects of stroma on chemosensitivity appears to be much lower, and the generalizability of these effects is less clear, as modulation of cell signaling is less capable of impacting the cellular damage created by chemotherapy.

Despite the substantial and constantly advancing knowledge of specific molecular mechanisms mediating direct therapy resistance, this knowledge is yet to translate into the ability to improve clinical therapy outcomes. To a large extent, this lack of progress reflects the fact that chemotherapy sensitivity represents a complex, multifactorial phenomenon that integrates a combined effect of multiple mechanisms acting at different scales (cell intrinsic, population, microenvironmental, systemic). Even at the scale of individual tumor cells, therapy resistance is not necessarily reducible to a single molecular mechanism (44,45). Further, chemotherapy responses cannot be fully reduced to the level of specific molecular mediators. For example, the efficiency of chemotherapies is limited by their ability to reach all tumor cells during the relatively short period of high drug concentration in circulation (46,47). However, the effects of these multiple mechanisms ultimately converge at the level of proliferation and death dynamics. Cancer is eradicated when cell death exceeds proliferation for a sufficiently long time to eliminate/permanently arrest all the tumor cells or to bring the disease burden to a threshold where the few surviving cells can be eliminated by the immune response. If this is not achieved, the cancer relapses. Despite this fact, considerations of proliferation/death dynamics are typically neglected when interpreting experimental data or designing clinical therapies. This omission is especially problematic for cytotoxic therapies that are administered in cycles, where rounds of acute drug exposure are spaced by several weeks of recovery, which is needed to manage therapy-associated side effects. For example, the common drug sensitivity assays, including sensitivities to chemotherapeutic agents, involve a single readout 3-4 days after the start of the treatment, often without wash-off of the drug, over a range of drug concentrations. These assays quantify the number of viable cells (via proxy readouts, such as MTT), relative to the number of cells in vehicle-treated control, with drug sensitivity usually assessed in metrics such as dose sufficient to induce a two-fold reduction in viability. The common implicit assumption is that a single timepoint measurement of net cell viability is sufficient to predict the overall therapy efficiency. However, a lack of response detectable at this time point might hide a complete elimination over a longer timeframe for chemotherapeutic agents that shift the proliferation/death balance (**Fig. 4, Fig. S7**). Moreover, identical short-term responses can reflect a wide range of proliferation/death combinations that can lead to dramatic differences over a longer time frame, where higher proliferation enables quick recovery and fast amplification of fitness differences.

In addition to consideration of temporal proliferation/death dynamics, understanding of therapeutic responses in vivo might also require consideration of space. For both direct resistance mechanisms, mediated by paracrine factors produced by stroma, and for proliferation-mediated effects uncovered in this study, the effects should be limited to tumor cells within sufficient proximity to stromal niches. Therefore, tumor/stroma ratios and specific patterns of the spatial distribution of stroma and tumor cells are expected to have a substantial impact on therapeutic responses. Moreover, the spatial patterns often change with treatment, as effective therapy typically leads to an increase in stroma/tumor ratio, as the numbers of neoplastic cells are reduced while neoplastic cell death leads to enhanced wounding signals, triggering stromal reaction/activation, *i.e*., enhanced secretion of inflammation-associated cytokines and increased ECM production, manifesting as swelling of stromal regions. While the diagnostic value of stromal activation is unclear (48), given the therapy-induced tumor shrinkage, it should increase the proportion of tumor cells within the spatial niche, where tumor cells are influenced by direct or proliferation-mediated resistance, thereby enhancing the importance of microenvironmental resistance mechanisms.

While our study focuses on enhanced proliferation within the peristomal niche, the importance of proliferation/death rates extends beyond the tumor-stroma crosstalk. Accelerated recovery between chemotherapy cycles might also be contributing to the stroma-independent link between a high proliferative index and poor outcomes. Both TNBC, which tend to be stroma-rich, and Ewing sarcomas, which tend to be relatively stroma-poor (49) tend to be highly sensitive to standard-of-care chemotherapies, yet these tumors typically resist therapeutic elimination. Based on strong experimental evidence, this discrepancy is often attributed to intratumor heterogeneity, enabling selection of intrinsically chemoresistant populations, or dormancy, i.e., some tumor cells avoid elimination either due to drug insensitivity or the ability to “sleep through the storm” through quiescence. Our results suggest that the high proliferation of residual tumor cells might also be an important contributor, suggesting parallels with r/K reproductive strategies in evolutionary ecology (50). Cell-intrinsic and cell-extrinsic mechanisms involving direct protection against the cytotoxic and cytostatic effects of the drugs are analogous to the K strategy, focused on enhanced protection/survival. In contrast, the indirect protection mechanism suggested by our study is analogous to the r strategy, where reproductive success is achieved by maximizing the number of offspring.

To some extent, the neglect of consideration of spatiotemporal proliferation/death dynamics reflects the lack of established experimental and analysis pipelines for accurate quantitative inferences of parameter values. Moreover, linear logical inferences used in molecular oncology studies are not suitable for understanding responses that integrate the impact of multiple variables. This task requires the use of biologically driven and parameterized mathematical modeling tools. The use of in silico modeling enables the understanding of the contribution of individual variables governing proliferation/death dynamics on therapy outcomes within biologically and clinically relevant parameter values, uncovering their emergent properties. The model’s simulations allow the exploration of large parameter space while uncoupling the studied parameters from confounding factors. The “clean” in silico experiments can provide full control over variables while addressing questions that are not accessible to purely experimental studies. For example, within purely experimental approaches, it is impossible to address whether a 5% difference in cell proliferation in stroma-proximal tumor cells can impact therapy outcomes (**Fig. 5G**). On the other hand, it is not possible to incorporate consideration of all factors shaping chemosensitivity. In addition to increasing the model’s complexity, the inclusion of additional variables is limited by the availability of knowledge on the additional parameters, as well as the ability to extract relevant quantitative information required for parameterization. Thus, inclusion of additional variables would necessitate the use semi-arbitrary assumptions, which reduces the utility of the model’s inferences. For example, our ABM does not include variabilities in cell-intrinsic chemotherapy sensitivities, differences in replication potential *etc*., as we lack the ability to extract relevant information from experimental and clinical samples. Therefore, the quantitative inference of modeling studies should be taken with caution, and their incorporation into the body of biological knowledge and clinical decision-making requires rigorous validation and follow-up.

In summary, our study uncovers a novel indirect chemoresistance mechanism where enhanced proliferation of tumor cells within peristomal niches facilitates tumor recovery between chemotherapy cycles, thus reducing the odds of therapeutic extinction for tumors at the brink of elimination (**Figs. 3A and 4G**). This mechanism might be partially responsible for the known link between high stromal content and poor therapeutic responses, although the link likely involves additional mechanisms, such as reduced drug penetration, induction of EMT/stemness, etc. Our study highlights the limitations of standard viability assays to understand the impacts of therapies while pointing out to the utility of understanding the spatiotemporal birth/death dynamics of cancer cells through the integration of experimental and mathematical modeling.

## Methods

### IHC staining

Formalin-fixed, paraffin-embedded tumors were cut at 5 microns sections. Deparaffinized tissue slices were blocked in PBS with 10% goat serum for 30 min at room temperature, then incubated at room temperature for 1 hour with primary anti-BrdU antibodies (1:100, Sigma#11170376001) and 1 hour with secondary antibodies (1:100, Vector Labs BA-9200), with 3×10 min washes after each incubation. The staining was developed using Vectastain ABC HRP Kit (Vector Labs, PK-6100), following manufacturer’s protocol. Cytoseal XYL mounting media (Epredia) was used to mount the slides.

### Histology image segmentation

IHC images were scanned with Aperio ScanScope XT Slide Scanner (Leica). The image areas corresponding to tumor tissue were segmented into BrdU+/-tumor cells, stroma, and necrotic tissue, with necrotic tissue excluded from the analyses (**Fig. S3A**). The segmentation was performed using an artificial intelligence (AI) based semi-automated deep learning quantification method (Aiforia version 5.2, Aiforia Inc, Cambridge, MA). Our training set for segmentation consisted of 20 slides, including samples after treatment and before treatment. An artificial intelligence-based software (Aiforia) was trained to recognize the nuclei of proliferating (stained with the proliferation marker BrdU) and non-proliferating tumor cells and record the x-y coordinates of the center of the nuclei. The stromal areas were segmented as polygons without separating cellular and ECM components. Segmentation accuracy was examined and approved by American Board of Pathology certified pathologist. Segmentation was performed on selected ROI for the initial correlation analyses. For proximity to the nearest stroma and RDS analyses, all of the tumor tissue was subjected to segmentation. Segmentation masks were downloaded as rds. Files for downstream analyses.

### Spatial statistics analyses

For the correlation analyses between the fraction of BrdU+ cells and stromal content, ROIs of 0.9 mm diameter were semi-randomly sampled from the tumor tissue area, avoiding large necrotic areas. Each ROI was segmented using the Aiforia AI model described above. The stromal area was calculated as a fraction of the ROI area, excluding necrotic tissue. The correlation between stroma and BrdU+ cells was calculated in GraphPad Prism using the simple linear regression method.

Advanced spatial statistics analyses were performed using R 4.1.2. The point pattern extraction from the Aiforia-segmented images has been performed using a combination of “sf” and “Magick” packages. Spatial statistics analyses were performed using the “Spatstat “, “kSamples”, “sp” and “Goftest”) packages. Polygonal segmentation of tumor stroma was tiled/pixelated using R package “sf” and “magick” into squares with the length of an average tumor cell diameter (15 μm). X-y coordinates of the center of each square (stroma unit/pixel) were recorded and used for the subsequent analyses.

For the analyses of distances to the nearest stroma, for each marker positive and marker negative (BrdU or Ki67) tumor cell, the Euclidian distance to the nearest stroma pixel was determined, and the distance frequencies and cumulative density graphs were plotted. Cumulative density functions of the respective distributions were used for analyses of the statistical significance of differences between distances of marker positive and marker negative cells by the Kolgomorov-Smirnoff test.

The RDF compares the average density of points against complete spatial randomness (CSR) across different length scales. To calculate the RDF for the distribution of marker positive/negative cells relative to the stroma, an annulus of width *dr* and radius *r* is placed around each of the stroma pixels. The number of marker +/-tumor cell centers within each annulus was calculated and divided by the expected number of tumor cell centers that fall inside the annulus under CSR. This calculation was repeated for each of the stromal pixels, and then the average value over all points at a certain *r* was recorded. CSR was generated *in silico* by maintaining the position of tumor cells and stromal pixels, as well as numbers of marker positive/negative tumor cells, but randomly shuffling tumor cell labels (marker status). For each sample, we generated a series of 39 CSF distributions and calculated a CSR average and 95% envelope of confidence.

### Agent-Based Modeling

The on-lattice ABM was developed using the Hybrid Automata Library (HAL) Java Library platform (34). In our model, cells are positioned in a 100 cells x 100 cells grid; the initial distribution of tumor cells and stroma is based on the Aiforia-extracted mask of an IHC slide image representing 1500 μm x 1500 μmarea. Each cell on the grid can be occupied by a tumor cell, stroma or be empty. Tumor cells are represented by agents that can divide, die, and move, while stromal pixels remain static. Time is discrete, with each time step representing 8 hours. At any timestep, the tumor cells can divide if there is an empty grid space in their immediate vicinity, die, or move to an available grid point in their vicinity (see **Fig. 3C**). The probability of proliferation is dictated by the proximity to the stroma, with the effect extending to tumor cells within 3 cells from the stromal pixels (based on the x-value of the maximum of the RDF). The proliferation rate was extracted from the labeling index given by the BrdU staining, while the death rate was taken as the difference between the net growth obtained by fitting the volume growth dynamics of the tumor and the proliferation rate (see the SI materials for the conversion from volume to cell fraction conversion and other details). Both apoptosis and death due to treatment occur without spatial bias throughout the grid, while the proliferation is increased by 5% -f 40% excess within the three-cell diameter radius during treatment. Based on our observations, in between chemotherapy cycles, both the proliferation and death rates are set to 55% of the initial value (in absence of treatment). The chemotherapy was simulated as 4 cycles and two scenarios of its effects were considered. First, chemotherapy induced a high kill rate over four days, followed by seventeen days where the proliferation and death rates were 55% of their respective rates before the treatment. Second, chemotherapy induced a low kill rate over two weeks with seven days of proliferation/death rates reduced as mentioned in the first scenario. Parameter values are described in Table S1. Details on the model and derivation of parameters from experimental data are provided in the Supplementary Methods.

### Cell lines and tissue culture conditions

Breast cancer cell lines were obtained from the following sources: MDA-MB-231, HCC1937, HCC1806, BT549, MDA-MB-436, HCC70, HS578T from ATCC and SUM149PT from Dr. S. Ethier (University of Michigan, Ann Arbor, MI). Breast cancer and normal fibroblasts were isolated from patient samples as previously described using protocols approved by the Dana-Farber Harvard Cancer Center (DF/HCC) Institutional Review Board. Fibroblast and SUM149PT cells were cultured in 50/50 mixture of DMEM-F12/MEGM with supplements containing 5% FBS; all other cell lines were cultured in in ATCC-recommended media. Cell line identities were confirmed by STR analysis. All cell lines and CAF isolates were routinely tested for mycoplasma contamination.

Firefly luciferase expressing derivates of TNBC cell lines were obtained as described in (25). Gaussia-expressing cell variants used for 3D assays were derived by lentiviral expression of Gaussia, subcloned from pMCS-Gaussia vector into pLenti6.3/V5-DEST vector (ThermoFisher). Fluorescently labeled cells for time-lapse microscopy studies were derived from the corresponding cell lines using a custom lentiviral vector which provides a nuclear expression of mCherry or GFP, coupled with puromycin expression, as described in (26). CAF-conditioned media was collected from confluent 10 cm dishes of CAFs grown in the fibroblast media. CAF CM experiments were performed in 50/50 mixtures of cell line-specific media with either CAF CM or fibroblast media.

### Cell viability readouts

For the luminescence-based readouts, firefly luciferase-expressing cells were grown in clear bottom white opaque 96 well plates (Corning, 3610). 30 min before taking the reading, the medium was replaced with fresh medium containing 25 μg/ml d-luciferin (GoldBio LUCK-2G). For the luminescence-based readouts of 3D assays, cells were seeded in ultra-low attachment U-bottom 96 well plates (Corning, 4520) at 3×10^4^ cells per well. Gaussia luciferase levels were measured using the supernatant media, after 5 hours of incubation with fresh media to avoid accumulation. 100 ul of 5 ug/ml Coelenterazine (GoldBio CZ) was injected using the luminometer. The luminescence signal was determined using GloMax luminometer (Promega); the background from the cell-free wells was subtracted from each reading. For microscopy-based readouts, time-lapse videos were generated with IncuCyte live cell imaging system (Sartorius) using ZOOM 4x objective. Images were acquired in red and green fluorescent channels as well as visible light channel every 12 hours for 7-21 days. Propidium iodide was added to the culture medium at 0.5 μg/ml to label dead cells. For the viability assays focused on assessing therapy responses, cells were seeded at 3.5×10^4^ cells per well to achieve ∼80% initial confluency. For the proliferation assays assessing the impact of CAFs and CAF CM on cell growth in the absence of treatment, cells were seeded at 3×10^3^ to achieve ∼5% initial confluency. For the fibroblast co-culture experiments, cells were plated in 3:1 TNBC: CAF ratios in 96 well-clear bottom tissue culture plates (Falcon #353072).

### Mouse xenograft studies

Tumors were initiated by orthotopic injection of 8 weeks old female NSG mice 1×10^6^ cells, suspended in 50/50 mix of DMEME-F12 culture media and Matrigel (BD Biosciences), in 100 μl volume; each animal was implanted with two tumors on the opposite flanks. Tumor growth was monitored weekly by electronic caliper measurements as described in (44). 30-40 min prior to euthanasia, mice were injected with 10 mg/ml BrdU dissolved in PBS. Tumors were fixed in formaldehyde and embedded in paraffin. All animal experiments were performed in accordance with the approved procedures of the IACUC protocol #IS00005557 of the H. Lee Moffitt Cancer Center.

### Clinical Samples

Diagnostic needle biopsy and post-surgery tissues of TNBC patients after AT-C neoadjuvant therapy were collected at Moffitt Cancer Center (Tampa, FL). All tissues utilized for this study were collected as part of Moffitt Cancer Center’s Total Cancer Care protocol (MCC#14690), with patients providing written informed consent. De-identified formalin-fixed paraffin-embedded breast tissues were cut at 5 microns sections and released in support of this study under IRB-approved protocol.

### Statistical analyses

Statistical analyses of *in vitro* and *in vivo* experimental data were performed using GraphPad Prism software, using statistical tests indicated in figure legends. Statistical analyses for the *in silico* simulations were performed with R and MatLab.

## Supporting information

Math Supplement

Siuppl video 4

## Data and code availability

All data supporting the findings of this study are available in the article and its Supplementary Information files. Tissue segmentation data is available at https://github.com/Marusyk-Lab/TNBC-CAF. The code for running the simulations and topology analysis is available online at https://github.com/ttanya86/TNBC

## Acknowledgments

This project is supported by the Florida Breast Cancer Foundation (A. Marusyk and D. Basanta), Moffitt Cancer Center Evolutionary Therapy Center of Excellence (A. Marusyk, D. Basanta and P.M. Altrock), William G.’Bill’ Bankhead Jr and David Coley Cancer Research Program 20B06 (A. Marusyk, D. Basanta and P.A. Altrock); S. G. Komen Research & Training grant (A. Marusyk), Richard O. Jacobson Foundation (P.M. Altrock), R37 CA244613 (Marusyk). This work has been supported in part by the Analytic Microscopy Core, Vivarium Core and the Tissue Core at the H. Lee Moffitt Cancer Center & Research Institute, a comprehensive cancer center designated by the National Cancer Institute and funded in part by Moffitt’s Cancer Center Support Grant (P30-CA076292). We thank the Moffitt SPARK program for supporting an internship for M.A.L. We would like to thank Jacob Scott and Bina Desai for their thoughtful comments and suggestions.

## References

1. Ge W, Yue M, Wang Y, Wang Y, Xue S, Shentu D, et al. A Novel Molecular Signature of Cancer-Associated Fibroblasts Predicts Prognosis and Immunotherapy Response in Pancreatic Cancer. Int J Mol Sci [Internet]. 2022 Dec 21 [cited 2023 Jan 12];24(1):156. Available from: https://pubmed.ncbi.nlm.nih.gov/36613599/

2. Xu J, Fang Y, Chen K, Li S, Tang S, Ren Y, et al. Single-Cell RNA Sequencing Reveals the Tissue Architecture in Human High-Grade Serous Ovarian Cancer. Clinical Cancer Research [Internet]. 2022 Aug 15 [cited 2023 Jan 12];28(16):3590–602. Available from: https://aacrjournals.org/clincancerres/article/28/16/3590/707396/Single-Cell-RNA-Sequencing-Reveals-the-Tissue

3. Irvine AF, Waise S, Green EW, Stuart B, Thomas GJ. Characterising cancer-associated fibroblast heterogeneity in non-small cell lung cancer: a systematic review and meta-analysis. Scientific Reports 2021 11:1 [Internet]. 2021 Feb 12 [cited 2023 Jan 12];11(1):1–15. Available from: https://www.nature.com/articles/s41598-021-81796-2

4. Jamieson NB, Carter CR, McKay CJ, Oien KA. Tissue biomarkers for prognosis in pancreatic ductal adenocarcinoma: a systematic review and meta-analysis. Clin Cancer Res [Internet]. 2011 May 15 [cited 2023 Jan 12];17(10):3316–31. Available from: https://pubmed.ncbi.nlm.nih.gov/21444679/

5. Graizel D, Zlotogorski-Hurvitz A, Tsesis I, Rosen E, Kedem R, Vered M. Oral cancer-associated fibroblasts predict poor survival: Systematic review and meta-analysis. Oral Dis [Internet]. 2020 May 1 [cited 2023 Jan 12];26(4):733–44. Available from: https://pubmed.ncbi.nlm.nih.gov/31179584/

6. Hu H, Piotrowska Z, Hare PJ, Chen H, Mulvey HE, Mayfield A, et al. Three subtypes of lung cancer fibroblasts define distinct therapeutic paradigms. Cancer Cell [Internet]. 2021 Nov 8 [cited 2023 Jan 12];39(11):1531–1547.e10. Available from: https://pubmed.ncbi.nlm.nih.gov/34624218/

7. Kramer CJH, Vangangelt KMH, van Pelt GW, Dekker TJA, Tollenaar RAEM, Mesker WE. The prognostic value of tumour-stroma ratio in primary breast cancer with special attention to triple-negative tumours: a review. Breast Cancer Res Treat [Internet]. 2019 Jan 15 [cited 2023 Jan 8];173(1):55–64. Available from: https://pubmed.ncbi.nlm.nih.gov/30302588/

8. Chakiryan NH, Kimmel GJ, Kim Y, Johnson JO, Clark N, Hajiran A, et al. Geospatial Cellular Distribution of Cancer-Associated Fibroblasts Significantly Impacts Clinical Outcomes in Metastatic Clear Cell Renal Cell Carcinoma. Cancers (Basel) [Internet]. 2021 Aug 1 [cited 2023 Jul 19];13(15). Available from: https://pubmed.ncbi.nlm.nih.gov/34359645/

9. Sahai E, Astsaturov I, Cukierman E, DeNardo DG, Egeblad M, Evans RM, et al. A framework for advancing our understanding of cancer-associated fibroblasts. Nat Rev Cancer [Internet]. 2020 Mar 1 [cited 2023 Jul 23];20(3):174–86. Available from: https://pubmed.ncbi.nlm.nih.gov/31980749/

10. Tao L, Huang G, Song H, Chen Y, Chen L. Cancer associated fibroblasts: An essential role in the tumor microenvironment. Oncol Lett [Internet]. 2017 [cited 2023 Jul 23];14(3):2611–20. Available from: https://pubmed.ncbi.nlm.nih.gov/28927027/

11. Cirri P, Chiarugi P. Cancer-associated-fibroblasts and tumour cells: a diabolic liaison driving cancer progression. Cancer Metastasis Rev [Internet]. 2012 Jun [cited 2023 Jul 23];31(1–2):195–208. Available from: https://pubmed.ncbi.nlm.nih.gov/22101652/

12. Farmer P, Bonnefoi H, Anderle P, Cameron D, Wirapati P, Becette V, et al. A stroma-related gene signature predicts resistance to neoadjuvant chemotherapy in breast cancer. Nat Med [Internet]. 2009 Jan [cited 2023 Jan 8];15(1):68–74. Available from: https://pubmed.ncbi.nlm.nih.gov/19122658/

13. Cleator S, Heller W, Coombes RC. Triple-negative breast cancer: therapeutic options. Lancet Oncol [Internet]. 2007 Mar [cited 2023 Jan 8];8(3):235–44. Available from: https://pubmed.ncbi.nlm.nih.gov/17329194/

14. Echeverria G V., Ge Z, Seth S, Zhang X, Jeter-Jones S, Zhou X, et al. Resistance to neoadjuvant chemotherapy in triple-negative breast cancer mediated by a reversible drug-tolerant state. Sci Transl Med [Internet]. 2019 Apr 17 [cited 2023 Jan 8];11(488). Available from: https://pubmed.ncbi.nlm.nih.gov/30996079/

15. Vangangelt KMH, Green AR, Heemskerk IMF, Cohen D, van Pelt GW, Sobral-Leite M, et al. The prognostic value of the tumor-stroma ratio is most discriminative in patients with grade III or triple-negative breast cancer. Int J Cancer. 2020 Apr 15;146(8):2296–304.

16. Östman A, Augsten M. Cancer-associated fibroblasts and tumor growth--bystanders turning into key players. Curr Opin Genet Dev [Internet]. 2009 Feb [cited 2023 Jul 23];19(1):67–73. Available from: https://pubmed.ncbi.nlm.nih.gov/19211240/

17. Takai K, Le A, Weaver VM, Werb Z. Targeting the cancer-associated fibroblasts as a treatment in triplenegative breast cancer. Oncotarget [Internet]. 2016 [cited 2023 Jul 23];7(50):82889–901. Available from: https://pubmed.ncbi.nlm.nih.gov/27756881/

18. Loh JJ, Li TW, Zhou L, Wong TL, Liu X, Ma VWS, et al. FSTL1 secreted by activated fibroblasts promotes hepatocellular carcinoma metastasis and stemness. Cancer Res [Internet]. 2021 Nov 15 [cited 2023 Jan 12];81(22):5692–705. Available from: https://aacrjournals.org/cancerres/article/81/22/5692/670494/FSTL1-Secreted-by-Activated-Fibroblasts-Promotes

19. Timperi E, Gueguen P, Molgora M, Magagna I, Kieffer Y, Lopez-Lastra S, et al. Lipid-Associated Macrophages Are Induced by Cancer-Associated Fibroblasts and Mediate Immune Suppression in Breast Cancer. Cancer Res [Internet]. 2022 Sep 15 [cited 2023 Jan 12];82(18):3291–306. Available from: https://aacrjournals.org/cancerres/article/82/18/3291/709018/Lipid-Associated-Macrophages-Are-Induced-by-Cancer

20. Raghavan KS, Francescone R, Franco-Barraza J, Gardiner JC, Vendramini-Costa DB, Luong T, et al. NetrinG1+ Cancer-Associated Fibroblasts Generate Unique Extracellular Vesicles that Support the Survival of Pancreatic Cancer Cells Under Nutritional Stress. Cancer Research Communications [Internet]. 2022 Sep 2 [cited 2023 Jan 12];2(9):1017–36. Available from: https://aacrjournals.org/cancerrescommun/article/2/9/1017/709220/NetrinG1-Cancer-Associated-Fibroblasts-Generate

21. Rosen S, Brisson BK, Durham AC, Munroe CM, McNeill CJ, Stefanovski D, et al. Intratumoral collagen signatures predict clinical outcomes in feline mammary carcinoma. PLoS One [Internet]. 2020 Aug 1 [cited 2023 Jan 12];15(8). Available from: https://pubmed.ncbi.nlm.nih.gov/32776970/

22. Győrffy B. Survival analysis across the entire transcriptome identifies biomarkers with the highest prognostic power in breast cancer. Comput Struct Biotechnol J. 2021 Jan 1;19:4101–9.

23. Ham IH, Lee D, Hur H. Cancer-Associated Fibroblast-Induced Resistance to Chemotherapy and Radiotherapy in Gastrointestinal Cancers. Cancers (Basel) [Internet]. 2021 Mar 1 [cited 2023 Jan 10];13(5):1–17. Available from: /pmc/articles/PMC7963167/

24. Hu Y, Yan C, Mu L, Huang K, Li X, Tao D, et al. Fibroblast-Derived Exosomes Contribute to Chemoresistance through Priming Cancer Stem Cells in Colorectal Cancer. PLoS One [Internet]. 2015 May 4 [cited 2023 Jan 10];10(5). Available from: /pmc/articles/PMC4418721/

25. Marusyk A, Tabassum DP, Janiszewska M, Place AE, Trinh A, Rozhok AI, et al. Spatial Proximity to Fibroblasts Impacts Molecular Features and Therapeutic Sensitivity of Breast Cancer Cells Influencing Clinical Outcomes. Cancer Res [Internet]. 2016 Nov 15 [cited 2023 Jan 10];76(22):6495–506. Available from: https://pubmed.ncbi.nlm.nih.gov/27671678/

26. Miroshnychenko D, Baratchart E, Ferrall-Fairbanks MC, Velde R vander Laurie MA, Bui MM, et al. Spontaneous cell fusions as a mechanism of parasexual recombination in tumour cell populations. Nat Ecol Evol [Internet]. 2021 Mar 1 [cited 2023 Jan 10];5(3):379–91. Available from: https://pubmed.ncbi.nlm.nih.gov/33462489/

27. Marusyk A, Tabassum DP, Altrock PM, Almendro V, Michor F, Polyak K. Non-cell-autonomous driving of tumour growth supports sub-clonal heterogeneity. Nature [Internet]. 2014 Oct 2 [cited 2023 Jul 20];514(7520):54–8. Available from: https://pubmed.ncbi.nlm.nih.gov/25079331/

28. Law R, Illian J, Burslem DFRP, Gratzer G, Gunatilleke CVS, Gunatilleke IAUN. Ecological information from spatial patterns of plants: insights from point process theory. Journal of Ecology. 2009 Jul;97(4):616–28.

29. Bull JA, Macklin PS, Quaiser T, Braun F, Waters SL, Pugh CW, et al. Combining multiple spatial statistics enhances the description of immune cell localisation within tumours. Sci Rep. 2020 Oct 29;10(1):18624.

30. Younge K, Johnston B, Christenson C, Bohara A, Jacobson J, Butler NM, et al. The use of radial distribution and pair-correlation functions to analyze and describe biological aggregations. Limnol Oceanogr Methods. 2006 Nov;4(10):382–91.

31. Brunton L, Lazo JS, Parker K, Buxton I, Blumenthal D. Goodman and Gilman’s Pharmacological Basis of Therapeutics. 11th ed. Mcgraw-Hill Professional; 2005.

32. Martín M, Seguí MA, Antón A, Ruiz A, Ramos M, Adrover E, et al. Adjuvant Docetaxel for High-Risk, Node-Negative Breast Cancer. New England Journal of Medicine [Internet]. 2010 Dec 2 [cited 2023 Jan 12];363(23):2200–10. Available from: https://www.nejm.org/doi/full/10.1056/nejmoa0910320

33. Chabner BA, Longo DL. Cancer Chemotherapy and Biotherapy: Principles and Practice (Chabner, Cancer Chemotherapy and Biotherapy). 5th ed. Lippincott Williams & Wilkins Next page; 2011. 1–812 p.

34. Bravo RR, Baratchart E, West J, Schenck RO, Miller AK, Gallaher J, et al. Hybrid Automata Library: A flexible platform for hybrid modeling with real-time visualization. PLoS Comput Biol [Internet]. 2020 [cited 2023 Jan 12];16(3):e1007635. Available from: https://journals.plos.org/ploscompbiol/article?id=10.1371/journal.pcbi.1007635

35. Nowakowski RS, Lewin SB, Miller MW. Bromodeoxyuridine immunohistochemical determination of the lengths of the cell cycle and the DNA-synthetic phase for an anatomically defined population. J Neurocytol [Internet]. 1989 Jun [cited 2023 Jan 10];18(3):311–8. Available from: https://pubmed.ncbi.nlm.nih.gov/2746304/

36. Gerlee P, Altrock PM. Extinction rates in tumour public goods games. J R Soc Interface [Internet]. 2017 Sep 1 [cited 2023 Jan 10];14(134). Available from: https://pubmed.ncbi.nlm.nih.gov/28954847/

37. Mross K, Maessen P, van der Vijgh WJ, Gall H, Boven E, Pinedo HM. Pharmacokinetics and metabolism of epidoxorubicin and doxorubicin in humans. Journal of Clinical Oncology. 1988 Mar;6(3):517–26.

38. Francescangeli F, De Angelis ML, Rossi R, Cuccu A, Giuliani A, De Maria R, et al. Dormancy, stemness, and therapy resistance: interconnected players in cancer evolution. Cancer Metastasis Rev [Internet]. 2023 Mar 1 [cited 2023 Jul 23];42(1):197–215. Available from: https://pubmed.ncbi.nlm.nih.gov/36757577/

39. Ishibashi M, Neri S, Hashimoto H, Miyashita T, Yoshida T, Nakamura Y, et al. CD200-positive cancer associated fibroblasts augment the sensitivity of Epidermal Growth Factor Receptor mutation-positive lung adenocarcinomas to EGFR Tyrosine kinase inhibitors. Scientific Reports 2017 7:1 [Internet]. 2017 Apr 21 [cited 2023 Jan 10];7(1):1–13. Available from: https://www.nature.com/articles/srep46662

40. Giannoni E, Bianchini F, Masieri L, Serni S, Torre E, Calorini L, et al. Reciprocal activation of prostate cancer cells and cancer-associated fibroblasts stimulates epithelial-mesenchymal transition and cancer stemness. Cancer Res [Internet]. 2010 Sep 1 [cited 2023 Jan 10];70(17):6945–56. Available from: https://aacrjournals.org/cancerres/article/70/17/6945/559773/Reciprocal-Activation-of-Prostate-Cancer-Cells-and

41. Wang W, Li Q, Yamada T, Matsumoto K, Matsumoto I, Oda M, et al. Crosstalk to stromal fibroblasts induces resistance of lung cancer to epidermal growth factor receptor tyrosine kinase inhibitors. Clinical Cancer Research [Internet]. 2009 Nov 1 [cited 2023 Jan 10];15(21):6630–8. Available from: https://aacrjournals.org/clincancerres/article/15/21/6630/75028/Crosstalk-to-Stromal-Fibroblasts-Induces

42. Hisamitsu S, Miyashita T, Hashimoto H, Neri S, Sugano M, Nakamura H, et al. Interaction between cancer cells and cancer-associated fibroblasts after cisplatin treatment promotes cancer cell regrowth. Hum Cell [Internet]. 2019 Oct 1 [cited 2023 Jan 10];32(4):453–64. Available from: https://link.springer.com/article/10.1007/s13577-019-00275-z

43. Ishii T, Suzuki A, Kuwata T, Hisamitsu S, Hashimoto H, Ohara Y, et al. Drug-exposed cancer-associated fibroblasts facilitate gastric cancer cell progression following chemotherapy. Gastric Cancer [Internet]. 2021 Jul 1 [cited 2023 Jan 10];24(4):810–22. Available from: https://link.springer.com/article/10.1007/s10120-021-01174-9

44. Vander Velde R, Yoon N, Marusyk V, Durmaz A, Dhawan A, Miroshnychenko D, et al. Resistance to targeted therapies as a multifactorial, gradual adaptation to inhibitor specific selective pressures. Nat Commun [Internet]. 2020 May 14 [cited 2023 Jan 10];11(1):2393. Available from: https://pubmed.ncbi.nlm.nih.gov/32409712/

45. França GS, Baron M, Pour M, King BR, Rao A, Misirlioglu S, et al. Drug-induced adaptation along a resistance continuum in cancer cells. bioRxiv [Internet]. 2022 Jun 25 [cited 2023 Jul 23];2022.06.21.496830. Available from: https://www.biorxiv.org/content/10.1101/2022.06.21.496830v1

46. Provenzano PP, Hingorani SR. Hyaluronan, fluid pressure, and stromal resistance in pancreas cancer. Br J Cancer [Internet]. 2013 Jan 15 [cited 2023 Jul 23];108(1):1–8. Available from: https://pubmed.ncbi.nlm.nih.gov/23299539/

47. Kim MJ, Gillies RJ, Rejniak KA. Current advances in mathematical modeling of anti-cancer drug penetration into tumor tissues. Front Oncol [Internet]. 2013 [cited 2023 Jul 23];3. Available from: https://pubmed.ncbi.nlm.nih.gov/24303366/

48. Fisher ER, Wang J, Bryant J, Fisher B, Mamounas E, Wolmark N. Pathobiology of preoperative chemotherapy: findings from the National Surgical Adjuvant Breast and Bowel (NSABP) protocol B-18. Cancer [Internet]. 2002 Aug 15 [cited 2023 Jul 20];95(4):681–95. Available from: https://pubmed.ncbi.nlm.nih.gov/12209710/

49. Tu J, Huo Z, Gingold J, Zhao R, Shen J, Lee DF. The Histogenesis of Ewing Sarcoma. Cancer Rep Rev [Internet]. 2017 [cited 2023 Jan 31];1(2). Available from: /pmc/articles/PMC5472389/

50. William D. Bowman SDH. Ecology. 5th ed. Sinauer Associates; 2020.

