## Supplementary Figures for "Paracrine enhancement of tumor cell proliferation provides indirect stroma-mediated chemoresistance via acceleration of tumor recovery between chemotherapy cycles"

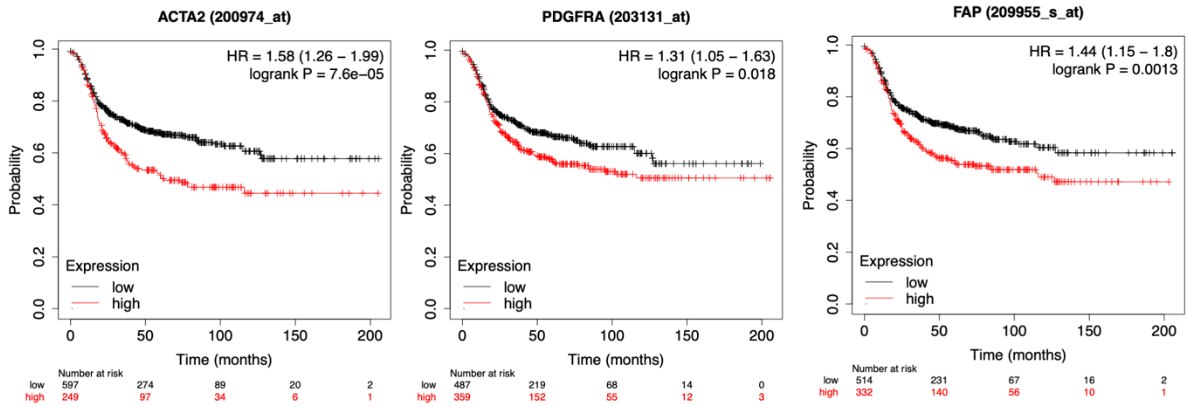


**Figure S1. Higher expression levels of CAF markers correlate with poor survival in TNBC patients**. Kaplan Mayer plots depict the analyses of the link between the expression of the indicated CAF markers and survival probability in TNBC patients. Analyses were performed with the KMplotter tool.

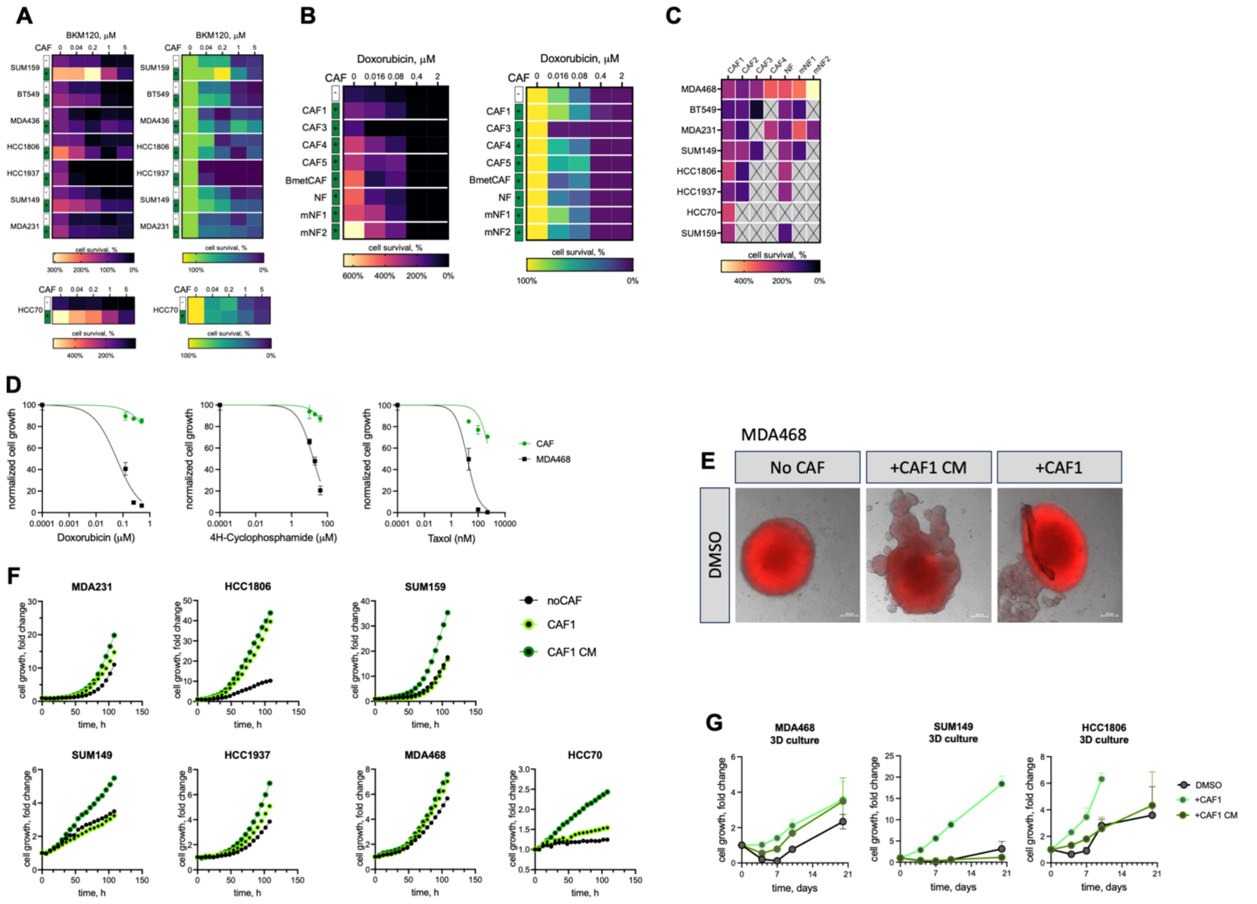


**Figure S2. Fibroblasts accelerate TNBC proliferation but do not provide direct chemoresistance. A** Heatmap summary of the impact of CAF co-cultures on BKM20 sensitivity of the indicated TNBC cell lines. **B, C** – cell survival rates for MDA468 cells in co-culture with a panel of CAFs of different origin (**B**), Heatmap summary of the impact of a panel of fibroblasts on doxorubicin sensitivity of MDA468 cell lines and CAFs of different origin (**C**). Heatmap summary on the impact of a panel of fibroblasts on baseline growth of the indicated TNBC cells. Normalization and color-coding scheme are depicted in Fig. 1B CAF1-5 denote CAF isolated from human breast cancers (CAF1 corresponds to the CAF isolate used in the main figures); BmetCAF denotes CAFs isolated from breast cancer brain metastasis; NFs denote fibroblasts isolated from normal breast tissue, mNF1,2 denote fibroblasts isolated from normal murine tissue (embryonic and lung respectively). **D** Sensitivity of CAFs to chemotherapy drugs compared to MDA468 cells. **E** Representative images of MDA468 organoids growing in CAF co-culture, with CAF CM or in regular medium 3 weeks after treatment with doxorubicin. MDA468 cells express mCherry marker. Scale bar is 200 µm. **F** Impact of CAFs and CAF CM on the baseline (no drug) growth of the indicated TNBC cell lines in 2D (**F**) and 3D (**G**) culture. Changes in cell numbers were monitored by time-lapse microscopy (**F**) and by luciferase expression (**G**).

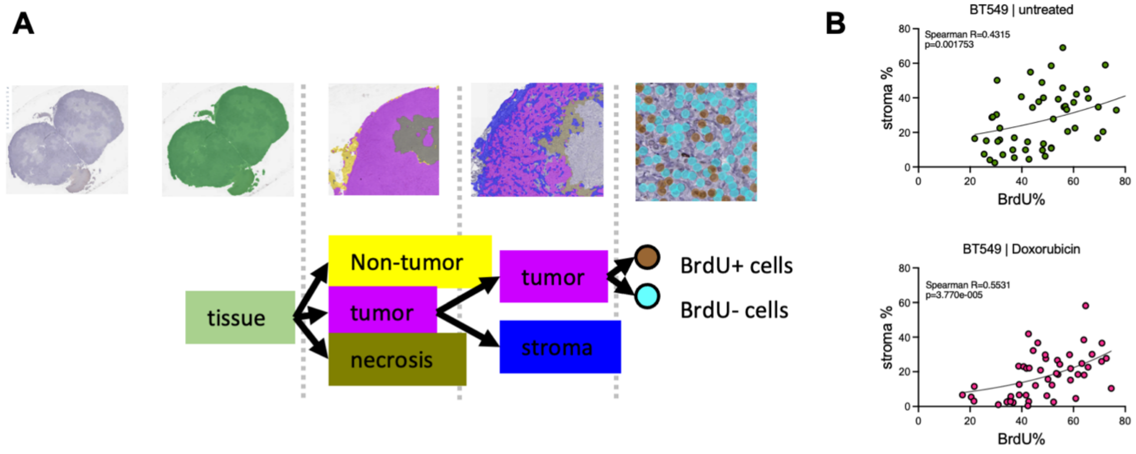


**Figure S3. Histological analyses of the link between stroma proximity and TNBC proliferation**. **A** Schemata of the histological image segmentation strategy. **B** Correlation between stromal content and proportion of BrdU+ cells in BT549 xenograft treated with vehicle or 2.5 mg/kg doxorubicin. Each data point represents an individual ROI as in Fig. 2A.

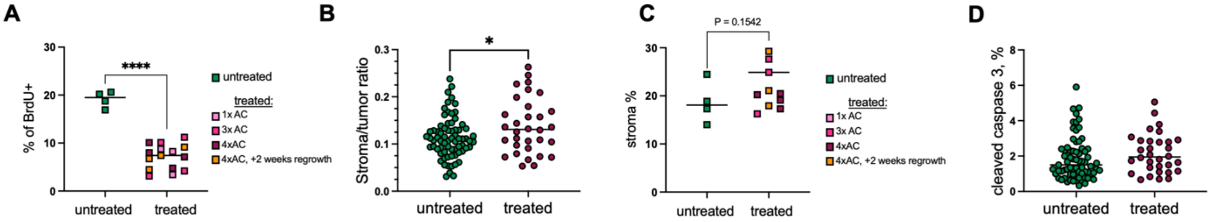


**Figure S4. Characterization of the impact of AC therapy**. Impact of AC therapy on the proportion of BrdU+ tumor cells (**A**) and stroma content (**C**) in MDA468 xenograft tumors treated with the indicated numbers of AC cycles (as in Fig. 3G). P value denotes the results of unpaired t-test. Each data point represents cross section of one xenograft tumor. Stroma/tumor ratio (**B**) and proportion of cleaved caspase 3+ cells the control and AC treated (4 cycles) tumors (**D**). Each data point represents an individual ROI as in Fig. 2A

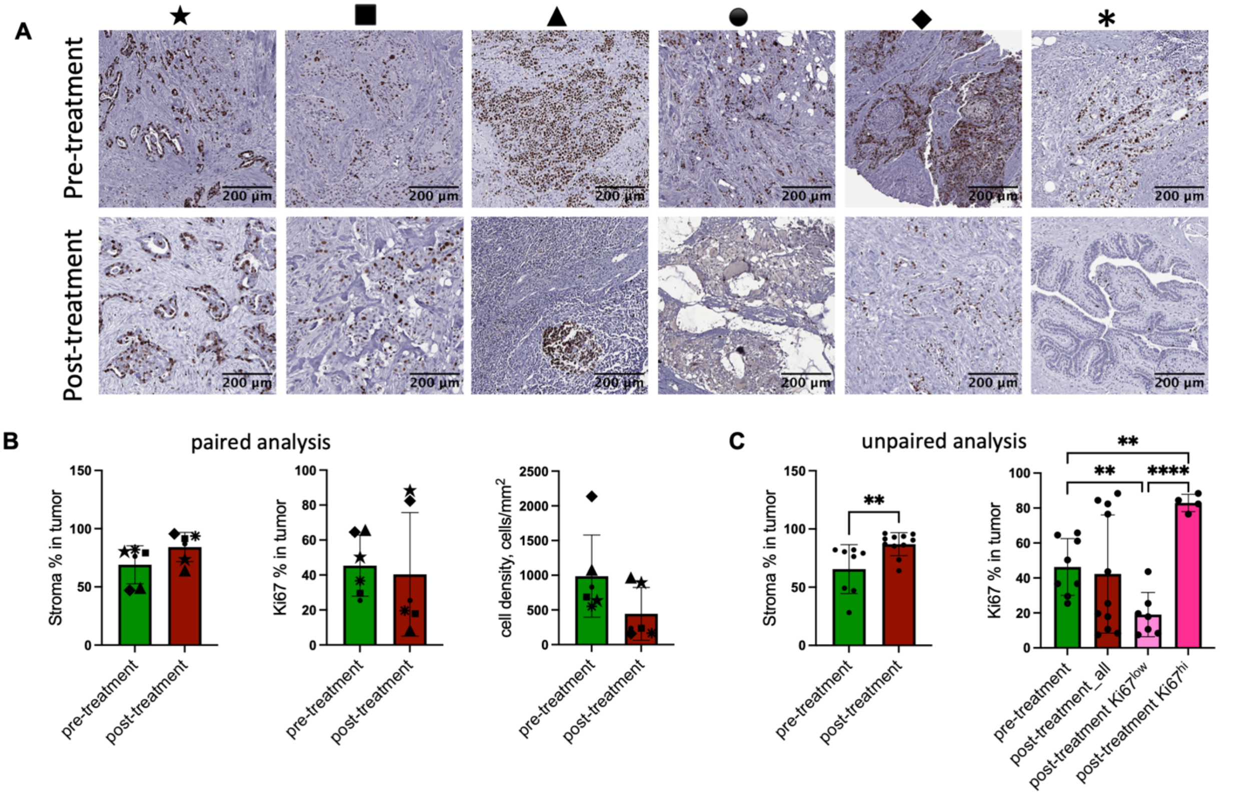


**Figure S5 Analysis of stroma content and cell proliferation in pre- and post-treatment primary TNBC tumors**. **A** Representative images of pre- and post-treatment tissue sections. **B** Human tissue samples were IHC stained for Ki67 expression. Segmentation of whole slide was performed by Aiforia algorithm. Stroma percentage is calculated for sromal&epithelial tissue excluding necrotic and adipose areas of the tissue samples. Ki67 percentage is % of Ki67+ positive cells from total number of cells detected in the tissue sample. For 6 patients pre- and post-treatments samples were available, data for each patient is represented by different symbol in paired analysis (★●■▲♦∗). No statistically significant difference was detected by paired t-test. **C** All available tissue samples were combined in pre- and post- treatment groups (N=8 and N=10, correspondingly). Statistical significance was defined using unpaired t-test. Difference in Ki67 expression between pre- and post-treatment samples was significant only when post-treatment samples were split onto two groups with hi and low Ki67 expression (p=0.0033 for hi, p=0.0072 for low). The difference between this groups is statistically significant based on t-test (p<0.0001).

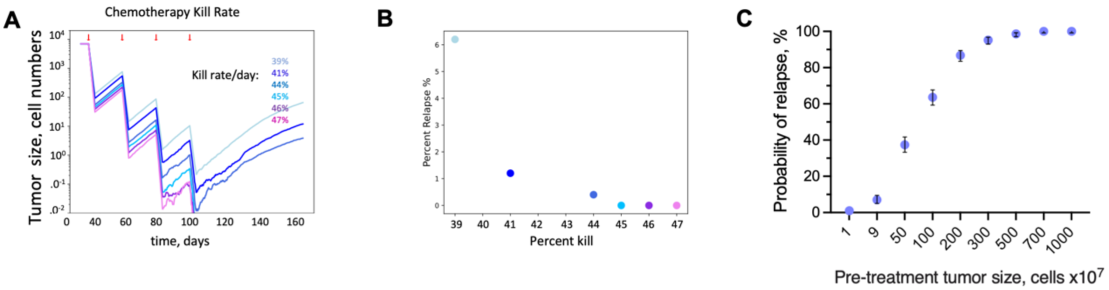


**Figure S6. In silico estimations of death rates.** The ABM model was used to estimate the death rates necessary to achieve complete penetration of tumor elimination in 500 in silico runs, given the experimentally observed net proliferation and birth rates of tumor cells. **A** The average number of tumor cells over time under different death rates. **B** Proportion of the tumors that survive the treatment in 500 in silico runs. **C** Dependence on sampling grid size of the tumor relapse for the simulations under short-term cytotoxic effects of the chemotherapy under 15% bias in proliferation due to stromal effects. A total of 10000 simulations have been run, and for each data point 500 samplings of 1,9,50, 100, 200, 300, 500 700 and 1000 simulations have been selected and verified for relapse. Each data point is the average of the 500 samplings with 95% confidence interval.

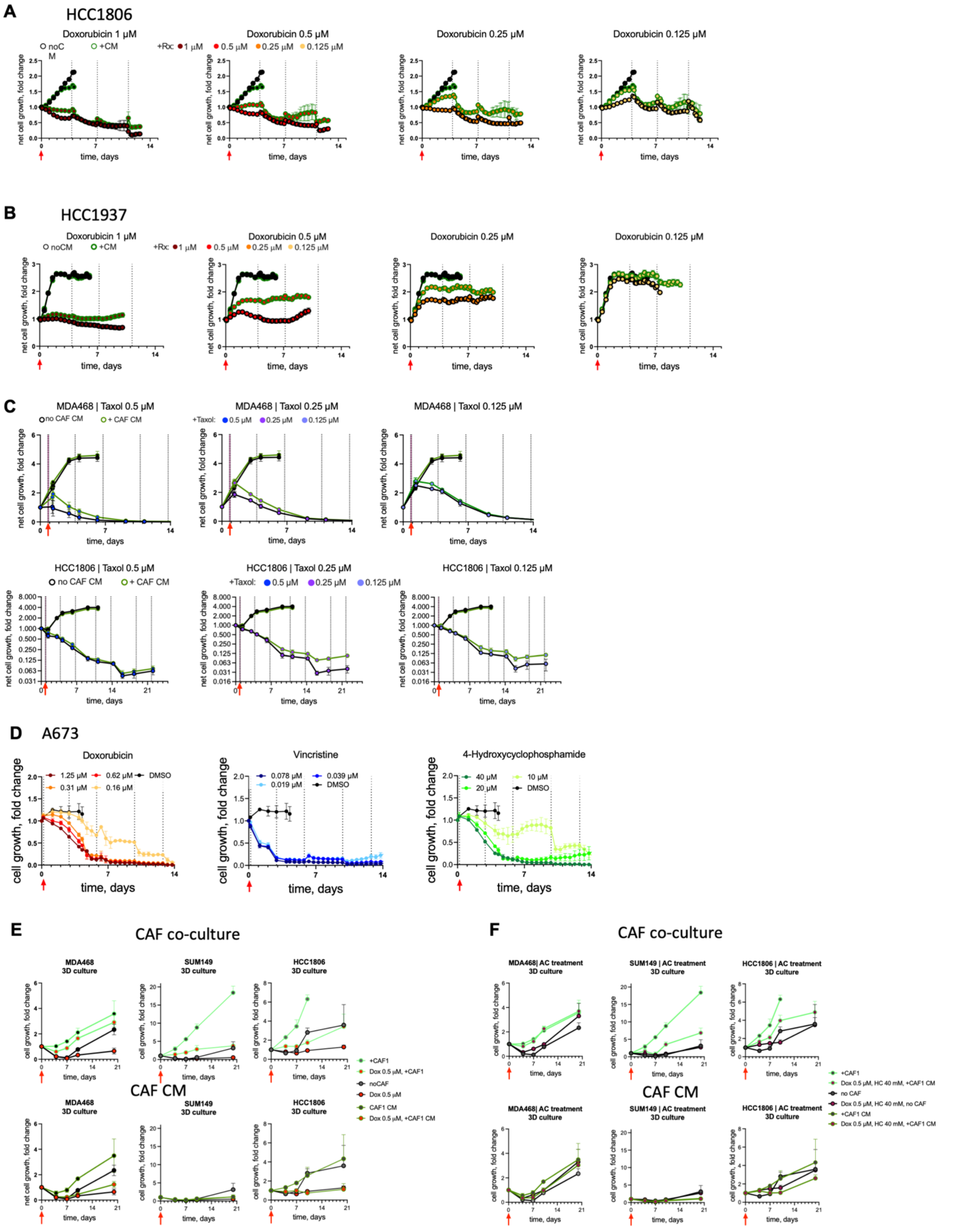


**Figure S7. Generalizability of the lasting increase in cell death following transient chemotherapy treatment. A, B** Net growth of TNBC cell lines HCC1806 (**A**) and HCC1937 (**B**) cells following the 2 hr exposure to the indicated concentrations of doxorubicin or DMSO vehicle control in the presence of control or CAF CM, assessed by time lapse microscopy. **C** Cell growth of MDA468 and HCC1806 following the 2 hr exposure to the indicated concentrations of Taxol or DMSO vehicle control in the presence of CAF CM. **D** Net growth of Ewing sarcoma cell line A673 following 6 hr exposure to the indicated chemotherapeutic agents. **E, F** Impact of CAFs and CAF CM on the growth of cell organoids after treatment with doxorubicin or combinational AC therapy (doxorubicin followed by 4-hydroxycyclophosphamide). Cell growth was monitored by luciferase expression.

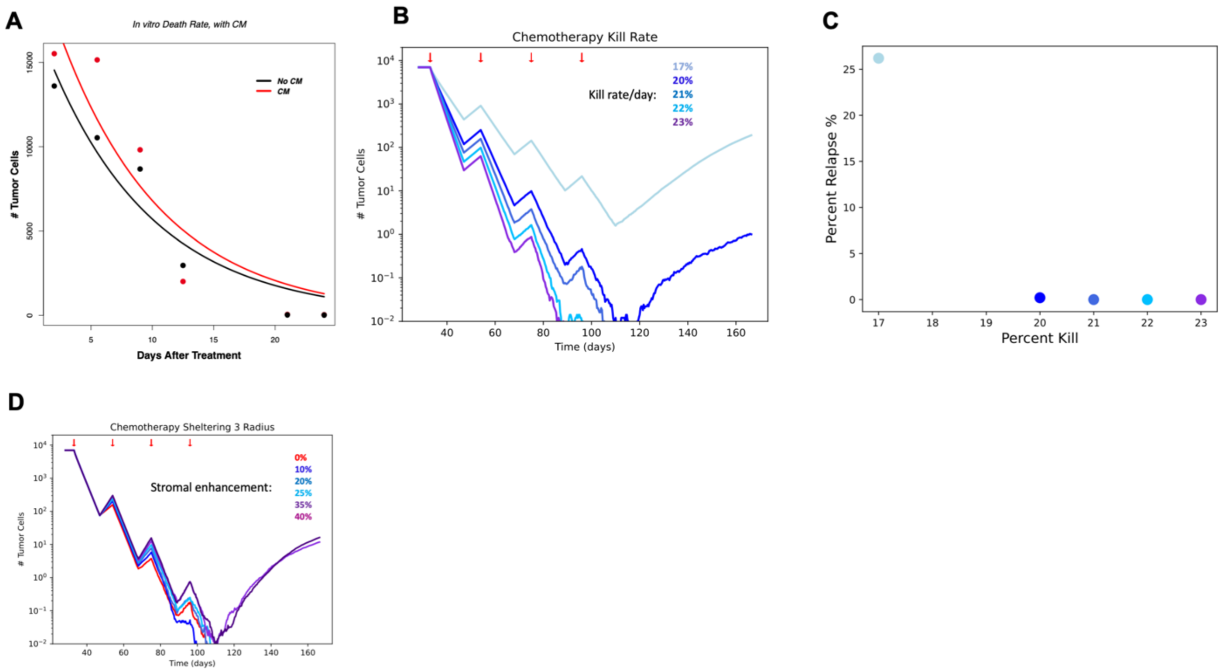


**Figure S8. Estimates of persisting death rates. A** Estimation of in vitro death rates of MDA468 cells with CAF CM and in control media following the 2 hr exposure to 0.5 μM doxorubicin. **B** Simulation of tumor cell population decline and growth during 4 cycles of chemotherapy. Different death rates parameters are explored. **C** Effect of various death rates on probability of tumor relapse. **D** Impact of the indicated magnitude of enhancement of cell proliferation within 3 cell diameters from stroma border on the average population size over the course of chemotherapy that, in the absence of stromal effects eliminates tumors with 0% relapse probability.

### Supplementary video:

**SV1** Representative run of an ABM simulation without stromal proliferation–enhancement effects.
**SV2** Representative run of an ABM simulation with stromal proliferation–enhancement effects.
**SV3** – Time lapse video of MDA468 cells response to a 6 hour exposure to 0.125 μM doxorubicin, in control media. Microscopy photos were taken every 12 hours.
**SV4** – Time lapse video of MDA468 cells response to a 6 hour exposure to 0.125 μM doxorubicin, in CAF CM. Microscopy photos were taken every 12 hours.
