## Supplementary material for "Paracrine enhancement of tumor cell proliferation provides indirect stroma-mediated chemoresistance via acceleration of tumor recovery between chemotherapy cycles": Math Supplement

### Spatial Analysis

We wanted to test the presence and quantify the observed bias in the localization of proliferating tumor cells in stroma's vicinity. We used a combination of artificial intelligence-based algorithm (Aiforia) and geometrical and spatial analysis packages in R to extract the point patterns of our histological samples and use them for spatial analysis (see Methods).

First, we looked at the distribution of the Euclidian distances between non-proliferating tumor cells to the nearest stroma cell clusters, and compared this distribution with the distribution of the distances from proliferating tumor cells to nearest stroma. We focused on three metrics. 1. the peak of the distributions indicating the range of the most frequent distances for either proliferating/non-proliferating tumor cells; 2. the median of the distribution, indicating the distance at which half of the (proliferating/non-proliferating) tumor cells would be encountered from the nearest stroma; 3. the standard deviation of the distribution, indicating the overall spread in the measured distances (with a higher spread indicating a tendency to a more uniform distribution of the distances).

We also calculated the cumulative density function (CDF) of the frequency distributions. These CDFs are direct read-out of the probability of finding a proliferating or non-proliferating tumor cell at a certain distance from stroma. Differences between CDFs was measured using the Kolgomorov-Smirnoff statistics (both distributions are poissonian in the median, and the standard deviations are typically close to each other).

As shown in Fig. 2, we calculated a positive stroma effect on proliferation in its vicinity for both the untreated samples as well as the samples collected after 4 cycles of chemotherapy. Specifically, for the untreated samples, both peaks (most likely values) and medians of the nearest-distance distributions of proliferating tumor cells occur at lower values than in non-proliferating tumor cells (20.05 $\mu m$ vs 23.624 $\mu m$ and 38.346 $\mu m$ vs 46.438 $\mu m$, respectively). Moreover, the standard deviation calculated from proliferating tumor cells is slightly lower than compared to non-proliferating tumor cells. These results indicate a tendency of proliferation occurring preferably closer to stroma, rather than being randomly distributed. At the same time, the CDFs for the proliferating and non-proliferating tumor cells led to the conclusion that proliferating tumor cells are typically found at closer stroma than non-proliferating tumor cells.

The radial distribution function (RDF, $g(r)$) approach (see Methods) also revealed that there is a 25% to 35% increase in proliferation within a three-cell diameter-distance from stroma (Fig. 3C), with a slowly decaying probability over a distance up to six cell diameters from away from stroma boundaries.

A similar stroma-tumor cell spatial interaction was observed in samples undergoing treatment, with increased intensity of the stromal effects on the proliferation. With a high consistency, we observed a similar phenomenon in an untreated human sample of TNBC stained with KI67 (Fig. 3D), although less pronounced (only around 5%). The fraction of all KI67-positive cells in these samples was typically close to 67% of the total cells. Thus, the amount of bias in the system can be limited by the availability of space in the stroma's vicinity. KI67-positive cells already occupy most of the space in the stroma's vicinity.

#### *ABM Parametrization*

Our goal was to develop a new agent-based modeling (ABM) approach to study scenarios in which stroma sheltering can prevent tumor elimination by chemotherapy and identify better treatment strategies. To this end, we used the customizable simulation platform HAL[1], and parametrized an ABM using an untreated sample to set the spatial stroma configuration and the growth dynamics. Thus, simulations of the spatially explicit system initially replicate the empirically observed settings as closely as possible.

The death rates of cancer cells were extracted from volume measurements and histological samples of untreated mice were estimated in the following way. Tumor (volume) growth in subcutaneous mouse xenograft models in the initial stages of growth can be described by an exponential growth law $V(t) = V_0 e^{rt}$, where *t* is time, *V(t)* is the volume of the tumor, $V_0$ is the initial detected volume of the tumor, *r* is the net growth rate which has two components, a proliferation rate and a death rate. Using the growth dynamics data and fitting an exponential growth law to that data (**see fig. S8**), we found a net positive growth rate of 0.04 $mm^3$ /day. Given that the number of tumor cells is directly proportional to the volume of the tumor through the cell volume density $\rho_{volume}$ ($N_{tumor\ cells} = V_{tumor} * \rho_{volume}$), the volume growth rate translates into a 4% net change in the cell count per day. Moreover, there is a linear relationship between the volume cell density and the area cell density, which we extract from histological samples, $N_V = \frac{N_A}{(w-d-2h)}$, [2]where $N_v$ is the volume cell density, $N_A$ is the area cell density, *w* is the histological sample thickness, *d* is the average cell diameter, and 2*h* is the spherical cap, which accounts for the loss of undetected parts of the cell in the histological sample and could be calculated as

$$h = \frac{d}{2-\sqrt{\frac{(d^2}{2)} - \frac{A_{crit}}{\pi}}},$$

where d is the average cell diameter, $A_{crit}$ is the minimum area detectable during the segmentation, typically ~10 $\mu m^2$. This calculation implies that the 4% net change in the tumor cell number can be extrapolated to a 4% net change observed in 2D histological samples. BrdU staining index in untreated samples is ~ 12% (BrdU staining detects nuclei in the S phase of the mitosis, which is approximately 8h, so around 36% of the tumor cells undergo division in a day), indicating that the death fraction is approximately 12 % - 4%/3 days=11% in three days. In our spatial simulation approach, the initial conditions were chosen to correspond to a tumor region away from the tumor boundary or from major necrotic areas. This mimics a tumor in dynamic equilibrium. Thus, the death rate dictates the maximum proliferation rate since proliferation can't occur when a cell's neighborhood is occupied (the ABM operates on contact inhibition). At the same time, the simulations are constrained by the fact that about 20% of the grid space has to be empty at all times to account for micronecrotic areas, which we implemented to better mimic the histological samples. Accounting for these constraints, and setting the proliferation fraction of the tumor cells in the histological samples, we arrived at a final death rate approximation of 0.125 cells/8h. Adjusting the proliferation rate to 0.2/8h, we obtain that 12.54% of the cells divide every 8h on average, whch we used in our simulations.

In the ABM approach, cell migration to a nearby available grid point was possible if, at that particular time, the cell did not divide or die. This cell movement approach recapitulates the relative movement due to the "pushing" of tumor cells (*in vivo*). Additionally, as can be seen from Fig. S4D, both in the untreated and the treated samples, apoptosis occurred with equal probability throughout the tumor tissue. Hence, we did not spatially bias the death rates in our model.

Treatment was assumed to reduce overall cell turnover. Based on our observations (Fig. S4A), during the chemotherapy and between chemotherapy cycles, the proliferation rate was set to 55% of the proliferation rate before treatment. Similarly, the death rate during the recuperation time in between the chemotherapy cycles, the death rate was set to 55% of the death rate before the treatment.

**4 Days High Cytotoxicity Scenario for Doxorubicin Mechanism**

Given both our *in vitro* and i*n vivo* results, we wanted to test whether stromal effects on proliferation in TNBC xenografts would lead to a faster recuperation of tumor cells numbers in between chemotherapy cycles, and whether this effect would lower the chances of tumor eradication after four cycles of treatment. We used the ABM approach parametrized as mentioned above and investigated the effects of chemotherapy both in presence and in absence of stromal effects. Our measurements of cell death/proliferation rates *in vivo* were taken 4 days after the $4^{th}$ (and last) cycle of chemotherapy (Doxorubicin) Hence, we initially simulated the chemotherapy to enhance the death rate of the tumor cells during 4 days, followed by reduced proliferation and death rates for 17 days (repeated 4 times to capture the 4 cycles of therapy).

To test the effectiveness of stroma to replenish the tumor in between therapy cycles, we first identified the kill rate necessary to eliminate the tumor in the absence of stroma (Fig. S4E-F), and at which values of stromal effects of up to a 3 cell-diameter distance the tumor would relapse after 4 cycles of therapy. We found that at rates equal and above 45%/8hours during 4 days, tumor cells could be completely eradicated. Under these killing rates, though, an increase in the proliferation rate for tumor cells within a distance of 45 μm from stroma by only 10%, lead to a 10% probability of tumor relapse (Fig. 3H-I). An increase in the proliferation rate by 25%, in the same manner, led to a 3% chance of relapse, despite eliminating 98.3% of the cells after the first cycle of chemotherapy. This shows that, indeed even a small increase in proliferation, potentially due to stroma, can lead to tumor escape from therapy.

**The total tumor cell count follows an exponential decay law during chemotherapy at high doses**

While we sampled over several doses in the *in vitro* experiments, only higher doses were used in our *in vivo* experiments. To understand the potential death dynamics *in vivo*, we focused on the death dynamics for the samples receiving higher doses *in vitro* (1 μm Doxorubicin). The respective experiments ran over more than 600 hours, and several growth media changes were needed, in between which the data indicated that there was little change in cell counts. We reduced that data so that a single average data point was taken for an interval between media changes, and we fitted the data with an exponential decay curve using least squares regression using "curve_fit" function from the the scipy.optimize package in Python.

As can be seen from Fig. S6A, at large doses of drug, there were no big differences in the cell counts dynamics, and the net growth rate (=proliferation rate – death rate) is close to -0.5%/h, measured over 2 weeks. Hence, the cell death at higher doses occurs over a longer period of time than previously considered and the cell population dynamics follows a slow exponential decay that has not been captured by our ABM. Given that we cannot directly integrate the *in vitro* results to parametrize an ABM that has been parametrized used *in vivo* data, we assumed that a similar death pattern is qualitatively recapitulated *in vivo*. We investigated 2 scenarios. First the cell death occurred with a high rate over 4 days followed by 10 days of a slightly increased death rate (hence, total 14 days of increased death rates; the rate of the lingering death was based on our observations the next day after the last treatment when we measure an apoptotic fraction increased around 10% from the baseline apoptosis rate observed in untreated samples). Second, death occurred according to an exponential law, with a lower constant rate compared with the high rate for the 4 days over 2 weeks. In our first scenario, we tested in our ABM approach the effects of an increased death rate by 10% at both 5% and 25% increased proliferation due to stroma. As shown in Fig. 3H-I, the excess death over 10 days after 4 days of high death rate (due to therapy), completely eliminated tumors after 4 cycles of chemotherapy if the excess proliferation due to stromal effects was below 5%. However, if the proliferation increase due to stroma was as small as 10%, the excess death over 10 days after each cycle was not enough to lead to tumor elimination.

We further assessed the effects of augmented proliferation in cells sufficiently close to stroma on tumor remission-relapse dynamics, given the scenario when death is exponentially decaying over 2 weeks. To this end, we used the same ABM approach as outlined above. First, we identified at with decay rate the tumor would be eliminated. Note here that even during chemotherapy, the proliferation rate is not zero, just reduced to 55%, as during the off-treatment time in between chemo cycles. Hence changes in the tumor cell counts always have contributions from proliferating and dying cells. As seen in Fig. S6B-C, we identified that a rate of 21%/8h or higher is needed to eliminate tumors after 4 cycles of Doxorubicin therapy (note that this is more than 2 times smaller than the rate needed to eliminate tumors if a high death rate would be present only over 4 days).

Next, using this lowest death rate that eliminates tumor cells consistently, we assessed how increased proliferation within 45 μm vicinity to stroma would affect the remission-relapse dynamics. In Fig. 4C-D it can be seen that an increase in the proliferation rate as high as 35% is needed to have a 0.5% chance of relapse. Interestingly, the tumor relapse rate after 4 cycles of chemotherapy in our simulations would not be further increased with higher proliferation rates. This may well be explained by a short period of recovery between cycles of therapy, as there likely is a low proliferation rate during the off-treatment.

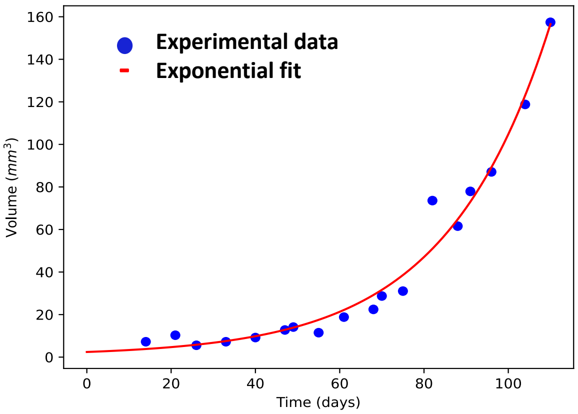


**Supplementary Figure 8**. Fitting tumor volume growth of an untreated mouse. Experimental data was collected using a caliper and a spherical geometry was assumed to calculate the volume of the tumor. Th experimental data was fitted with an exponential and the net growth rate was obtained from the fit.

### Supplementary Table 1. ABM parameters

| Parameter | Description | Value | Units | Source |
|---|---|---|---|---|
| ABM grid size | The 2D simulation grid used in the ABM | 100x100, corresponding to 1500 $\mu m$ x 1500 $\mu m$ | NA or 2250000 $\mu m^2$ | User defined/histological sample-based point pattern |
| $\Delta t$ | time step of the ABM | 8 | h | User defined, based on the reported values of the S phase of the cell cycle measured with BrdU/EdU[3,4] |
| $p_{p0}$ | Probability of a cell to divide within a time step in absence of stromal effects | 0.2 | NA | Based on BrdU LI and adjusted based on the observation of ~20% micronecrosis |
| $p_{pt}$ | Probability of a cell to divide within a time step in absence of stromal effects during/after treatment | 0.11 | NA | experiment based |
| $p_{d0}$ | Probability of a cell to die within a time step in absence of stromal effects | 0.125 | NA | Based on BrdU LI and adjusted based on the observation of ~20% micronecrosis, and net growth rate extracted from untreated tumor volume growth |
| $p_{dt}$ | Probability of a cell to die within a time step in absence of stromal effects after treatment | 0.074 | NA | experiment based |
| $r_3$ | Radius for stromal effects | 3 grid distances | NA/$\mu m$ | Based on RDF $g_{maxx}$ 5 |
| $p_{ps}$ | Probability of a cell to die within a time step in presence of stromal effects without treatment | $p_{p0}$ + $p_{p0}$* stromal percent bias | NA | |
| $p_{pst}$ | Probability of a cell to die within a time step in | $p_{pt}$ + $p_{pt}$* stromal percent bias | NA | |

| | | | | |
|---|---|---|---|---|
| | presence of stromal effects during/after treatment | | | |
| $p_{dst}$ | Probability of a cell to die within a time step in presence of stromal effects after treatment | 0.068 | NA | Experiment based |

References


1. Bravo, R. R. *et al.* Hybrid Automata Library: A flexible platform for hybrid modeling with real-time visualization. *PLOS Comput. Biol.* **16**, e1007635 (2020).
2. Mi, H. *et al.* Digital Pathology Analysis Quantifies Spatial Heterogeneity of CD3, CD4, CD8, CD20, and FoxP3 Immune Markers in Triple-Negative Breast Cancer. *Front. Physiol.* **11**, 583333 (2020).
3. Massey, A. J. Multiparametric Cell Cycle Analysis Using the Operetta High-Content Imager and Harmony Software with PhenoLOGIC. *PLOS ONE* **10**, e0134306 (2015).
4. Bialic, M., Al Ahmad Nachar, B., Koźlak, M., Coulon, V. & Schwob, E. Measuring S-Phase Duration from Asynchronous Cells Using Dual EdU-BrdU Pulse-Chase Labeling Flow Cytometry. *Genes* **13**, 408 (2022).
5. Bull, J. A. *et al.* Combining multiple spatial statistics enhances the description of immune cell localisation within tumours. *Sci. Rep.* **10**, 18624 (2020).
